# Integration of SYT1 Interactomics and Dual-Localization Proteomics Links ER-PM Contacts to Lignin Deposition

**DOI:** 10.64898/2026.07.06.736826

**Authors:** Jorge Morello-López, Ulises Galvan, Francisco Benitez-Fuente, Vedrana Marković, Harriet Parsons, Tim J. Stevens, Jessica Pérez-Sancho, Vitor Amorim-Silva, Jelle Van Leene, Geert de Jaeger, Lourdes Rubio, Yvon Jaillais, Miguel A. Botella

## Abstract

Membrane contact sites (MCSs) are evolutionarily conserved intracellular nanodomains that physically bridge opposing lipid bilayers to facilitate non-vesicular communication and maintain cellular homeostasis. In plants, endoplasmic reticulum-plasma membrane (ER-PM) contact sites play fundamental roles in environmental adaptation, and are populated by specialized proteins which act as tethers such as Synaptotagmin 1 (SYT1). However, a comprehensive view of the molecular machinery governing processes at these junctions is still needed. In this work, we integrate affinity purification mass spectrometry, TurboID proximity labeling, and a dual-localization reanalysis of HyperLOPIT spatial proteomics to functionally map the protein interaction landscape of the ER-PM contact sites protein SYT1. Beyond recovering established ER-PM MCS functions, our analysis identified uncharacterized proteins as bona fide resident components of these junctions, and revealed that these nanodomains act as docking platforms that anchor the monolignol biosynthetic complex. By spatially organizing Membrane Steroid Binding Proteins and cytochrome P450 enzymes, our findings support a model where SYT1-mediated anchoring of this metabolon to ER-PM contact sites optimizes monolignol export required for stress-induced lignification. Ultimately, this proteomic framework expands the functional repertoire of ER-PM contact sites, opening new avenues to uncover hidden roles of MCSs across diverse eukaryotic systems.

## INTRODUCTION

Membrane contact sites (MCSs) are specialized subcellular nanodomains present in all eukaryotes in which two membranes become closely apposed without undergoing fusion^1,2^. The distance between the two membranes at the MCS typically ranges from 10 to 30 nm and is maintained by tethering proteins that physically bridge the opposing lipid bilayers^3,4^. MCSs are now recognized as a conserved feature of eukaryotic cells and are present in virtually all organelles, where they facilitate non-vesicular exchange of lipids, metabolites, small molecules, and signaling cues, thereby contributing to coordinated organelle function and dynamic cellular communication^5–7^.

Among the different types of MCSs, contacts between the endoplasmic reticulum (ER) and the plasma membrane (PM), known as ER-PM contact sites, are particularly prominent^8–10^. These domains are found across plants, animals, and yeasts, and they participate in several fundamental processes, including non-vesicular lipid transport^11^, regulation of calcium homeostasis^12^, and maintenance of cortical ER architecture^13^.

A key group of proteins involved in ER-PM tethering in plants belong to the synaptotagmin (SYT) family, represented in animals and yeast by related extended synaptotagmins and tricalbins, respectively^14^. *Arabidopsis thaliana* SYT proteins share a conserved domain organization: a N-terminal transmembrane region that anchors the protein in the ER membrane, a Synaptotagmin-like Mitochondrial-lipid-binding Protein (SMP) domain, with lipid-binding activity and dimerization capacity^11,15–17^, and two C-terminal C2 domains that mediate interaction with negatively charged phospholipids at the PM, mainly phosphoinositides^17–20^.

There are now studies highlighting the importance of SYT1 in multiple biological processes. SYT1 plays a role in resistance to salt and cold stress^17,19^, freezing tolerance^21^, viral infection^22–24^, and immune responses^25^. At the cellular levels, SYT1 is required for the formation of the tubular ER network^13^ and maintenance of ER morphology^26^. Recently, it has been reported that SYT1 plays roles in mechanosensitive ion channel function^13^, diacylglycerol transport in response to stress^17,18^, and cell-to-cell connectivity via plasmodesmata during innate responses^27^. Furthermore, our companion manuscript also establishes a role for SYT1 in the regulation of polarized growth of root hairs during normal development (Marković et al., 2026).

Beyond the SYT family, plant ER-PM contact sites are also populated by other proteins that perform a tethering function, such as the VAP27 (VAMP-associated protein 27) and MCTP (Multiple C2 domain and Transmembrane region Protein) families^14^. VAP27 proteins are anchored to the ER via a C-terminal transmembrane domain and normally interact with the PM through protein adaptors, contributing to the stabilization of the cortical ER network^13,27,28^. MCTPs are characterized by multiple N-terminal C2 domains that dock to the PM and a C-terminal region that inserts into the ER membrane, playing key roles in cell signaling and intercellular communication^29,30^. Together, these diverse tethering systems cooperate with SYT proteins to support the functional versatility of the ER-PM interface^14^.

All this information supports the view that the function of SYT1 is not limited to tethering or lipid transport but also includes additional roles during plant development and stress responses. In this article, we combined affinity purification coupled to mass spectrometry with proximity labeling assays to uncover the SYT1 interaction landscape. By merging these proteomic approaches with protein spatial resolution data, we not only validated known SYT1 interactors but also discovered novel candidates that point to uncharacterized roles for ER-PM contact sites. Our findings are supported by the specific localization of several newly identified proteins to these nanodomains. Notably, we further uncovered an association between SYT1 and proteins involved in monolignol biosynthesis at ER-PM contact sites. Using genetic and biochemical analyses, we demonstrate that SYT1 is involved in stress-induced lignin deposition, thus reporting an uncharacterized specific function of ER-PM contact sites in plants. Overall, the data provided in this study advance current knowledge of ER-PM contact site function in plants and provide an integrative basis for further exploring the role of these nanodomains in other biological processes.

## RESULTS

### A dual-omics proteomic strategy maps a high confidence SYT1 interactome

To obtain a proteome-wide view of the molecular environment associated with ER-PM contact sites, we combined affinity purification coupled to mass spectrometry (AP–MS)^31^ with TurboID-based proximity labeling (PL)^32^ of SYT1. This dual strategy yields highly complementary datasets, as AP–MS allows the isolation of stable and robust physical complexes, whereas PL successfully captures the surrounding spatial neighborhood of the bait, mapping proteins in immediate physical proximity to SYT1 despite their transient nature or low binding affinity. As C-terminally tagged SYT1 variants have already been shown to maintain functionality of the protein^17^, the GSrhino and TurboID tags (for AP-MS and PL respectively) were fused to the C-terminus of SYT1^32^. To standardize our biological system and facilitate the comparison with the downstream LOPIT spatial proteomics platform, *Arabidopsis* suspension cell cultures were systematically used as the starting material for all proteomic approaches. For AP-MS, given that SYT1 is anchored to the endoplasmic reticulum (ER) via an N-terminal transmembrane domain, membrane protein extraction was optimized by supplementing the standard isolation buffer with 1% digitonin, followed by dithiobis(succinimidyl propionate) (DSP) crosslinking to stabilize proximal interactions during pull-down. For proximity labelling, the neighbor SYT1 interactors were biotinylated *in vivo* and subsequently enriched using streptavidin beads under denaturing conditions. To ensure a high-confidence dataset, candidates from both proteomic approaches were filtered by comparing the normalized spectral abundance factors (NSAFs) of each identified protein against corresponding large AP-MS and PL cell culture control dataset. The results were filtered with a stringent criterion of an Affinity Enrichment Score (AES), which integrates both NSAF-based fold changes and T-test p-values (see Material and Methods). The stablished threshold for both techniques was an AES ≥ 20.

The combination of AP-MS and PL identified a total of 289 proteins in the SYT1 interactome, of which 37 were shared between the two techniques, 143 were unique to AP-MS, and 109 were unique to PL. The lists of proteins identified by each individual technique, along with those shared between them, is provided in **Supplementary Data 1**.

### SYT1 is primarily associated with structural ER tethers and lipid-related functions

To obtain an integrative overview of the generated datasets, we constructed a protein-protein interaction (PPI) network of the identified SYT1 interactors. This network recovered numerous well-established tethering and membrane-shaping components, strongly validating biological relevance and quality of our dataset (**Fig. 1a**). To facilitate visualization, text labels are restricted to those nodes shared between the two proteomic datasets and to candidates with previously documented roles MCS (see **Supplementary Figure 1** for the network with all nodes labeled). Among SYT1 interactors, we identified proteins from all three canonical ER-PM contact sites tether families (SYTs, VAP27s and MCTPs)^14^, previously validated SYT1 partners^17,33^, and novel, uncharacterized proteins. The relation between SYT1 and all documented candidates is specified in **Supplementary Table 1.** The dataset exhibits a notable enrichment of Synaptotagmin family members (including the bait SYT1, as well as SYT4 and SYT5), and related SMP domain-containing proteins CLB1, NTMC2T6.2, and TEX2B. Furthermore, core members of the VAP27 family (VAP27-1, −2, −3, and -4) and MCTP family (MCTP3, -4, -15, and -16) also appeared as interactors. Simultaneously, the network is also populated by a broad set of proteins involved in establishing and maintaining ER morphology, such as the reticulons (RTNLB1, −2, −3, - 4, -5, -8, and RTN20), LUNAPARK2 (LNP2) and ROOT HAIR DEFECTIVE 3 (RHD3)^34–37^.

**Figure 1.**
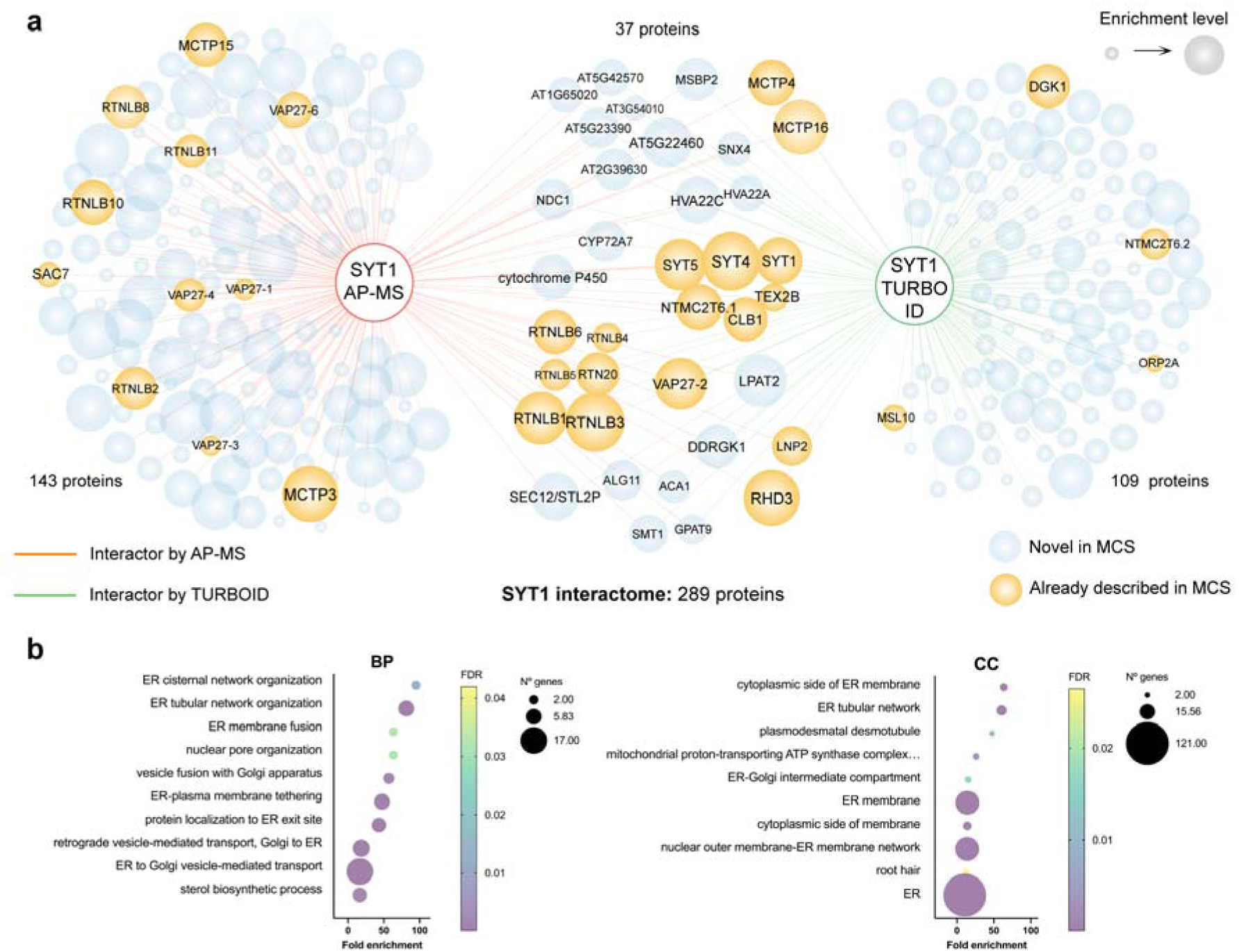
A dual-approach SYT1 interactome maps biological processes operating at ER-PM contact sites. **a, Combined protein-protein interaction network of SYT1 high-confidence interactors.** The network integrates datasets obtained from affinity purification mass spectrometry (AP-MS, red edges) and proximity labeling (TurboID, green edges). Nodes represent identified interacting proteins, color-coded to distinguish novel candidates (light blue) from components previously described in the literature as being related with ER-PM contact sites (orange, detailed in Supplementary Table 1). Node and label sizes are proportional to the log_10_ of a cumulative rank score, calculated by ranking proteins according to their enrichment values within each dataset and summing values for shared candidates. **b, Gene Ontology (GO) enrichment analysis of the global SYT1 interactome.** Dot plots display the top significantly enriched categories for Biological Process (BP, left) and Cellular Component (CC, right). X-axis indicates fold enrichment; bubble size corresponds to the number of genes assigned to each term, and the color gradient represents the false discovery rate (FDR) significance level.

The differential recovery of certain candidates across the techniques directly reflects the distinct biochemical natures of the two proteomic approaches. For instance, the capacity of TurboID PL to capture *in vivo* interactions that might be dynamic or transient is illustrated by the detection of diacylglycerol kinase 1 (DGK1), an ER-resident protein that specifically functions at MCS through its direct interaction with SYT1^18^. In contrast, several members of the VAP27 family were detected exclusively by AP-MS, which might be consistent with VAP27 proteins assembling into independent macromolecular complexes or establishing VAP27-specific microdomains adjacent to those organized by SYT1, without strictly residing within the same MCS^13^.

Gene Ontology enrichment analysis of the combined SYT1 interactome revealed a set of terms that align with SYT1 structural and physiological features. For instance, enriched Biological Process (BP) and Cellular Component (CC) terms explicitly grouped around ER network morphology, membrane remodeling, and active intracellular transport (**Fig. 1b**), functions that match established roles of MCS^13,26^. Indeed, many of these features correspond to well-established functional hallmarks of plant ER-PM contact sites, where structural maintenance is closely linked with lipid homeostasis, membrane trafficking dynamics, and cellular signaling^18,30,38,39^. Molecular Function (MF) analysis (**Supplementary Figure 2**) also highlighted the expected presence of steroid-related processes^40^ or lipid / phospholipid binding activities^19,41,42^. A clear example illustrating this last function is the phosphatase SAC7, a SYT1 interactor whose relation with MCS is characterized in detail in the companion manuscript (see Marković et al., 2026). However, other less anticipated functions also emerged from the analysis, such as monooxygenase activity, which is explored further below in the context of the ER-associated lignin metabolon.

### Re-analysis of LOPIT data identifies dual-localized proteins at the ER-PM interface and other boundaries

To systematically map components that may function at membrane junctions, we leveraged spatial proteomics data obtained via LOPIT (Localization of Organelle Proteins determined by Isotope Tagging)^43^. The LOPIT technique generates profiles of protein abundance along a density centrifugation gradient after gentle cellular lysis, such that proteins from the same subcellular organelle/compartment exhibit similar, characteristic gradient distributions^43,44^. A recent advancement in this technique has enabled dual-localized proteins not only to be identified but also to be assigned to specific organelle pairs, with a proportional measure of a protein population distribution between organelles^45,46^. Additionally, this advancement addresses missing values within and between replicates, enabling the addition of over 1000 proteins to spatial maps. Denoising protein gradient profiles has dramatically improved the resolution of these maps. Accordingly, this allowed us to reanalyze our existing experimental data to include subcellular localization probabilities for each protein^47^. This dual-localization data is publicly available and hosted as the interactive, open-access web application Choragraph (available at choragraph.org), which allows independent querying of subcellular protein distributions^47^.

Here, the underlying data were obtained from eight LOPIT experiments performed on Arabidopsis cell suspension cultures^45^, with two sample preparation methods (with and without carbonate washing) and combined into a single HyperLOPIT dataset containing 84 abundance values for 9441 proteins^47^. The Deep Neural Network (DNN) based analysis of proteomic data presented by Choragraph is specifically designed to identify proteins with LOPIT profiles that are best explained by a proportional mixture of different organelles. From the available outputs in the platform, we specifically used the data from Arabidopsis Comb v1 for downstream analyses. This enabled us to screen for candidate proteins showing high probability of localization to both the ER and other target organelle membranes, namely the PM, Golgi, trans-Golgi network (TGN), vacuole, chloroplast, or mitochondria.

As MCS-resident components are typically anchored to the ER membrane, we defined our ER-PM-localized proteome as proteins with Choragraph threshold proportions of ≥ 10% in the ER and ≥ 5% in the PM, allowing the identification of 324 dual-localized candidates (**Table 1**, **Supplementary Figure 3a**). The selected parameters recovered most members of the major canonical ER-PM tethering families, including SYT1, SYT5, CBL1, MCTP4, MCTP16, and the majority of the VAP27 family (VAP27-1, −2, and -6)^14^ (**Supplementary Figure 3b**). Next, we used the same allocation parameters used for dual ER-PM localized (≥ 10% for the ER and ≥ 5% for the respective target compartment) to comprehensively map the landscape of different ER–organelle interfaces across the HyperLOPIT dataset (**Table 1**). The complete LOPIT dataset and specific ER-organelle sub-lists are detailed in **Supplementary Data 2.** While detailed studies of specific organelles might require independent threshold refinement, this approach provides a first comparative framework for assessing global connectivity from the plant ER.

**Table 1.**
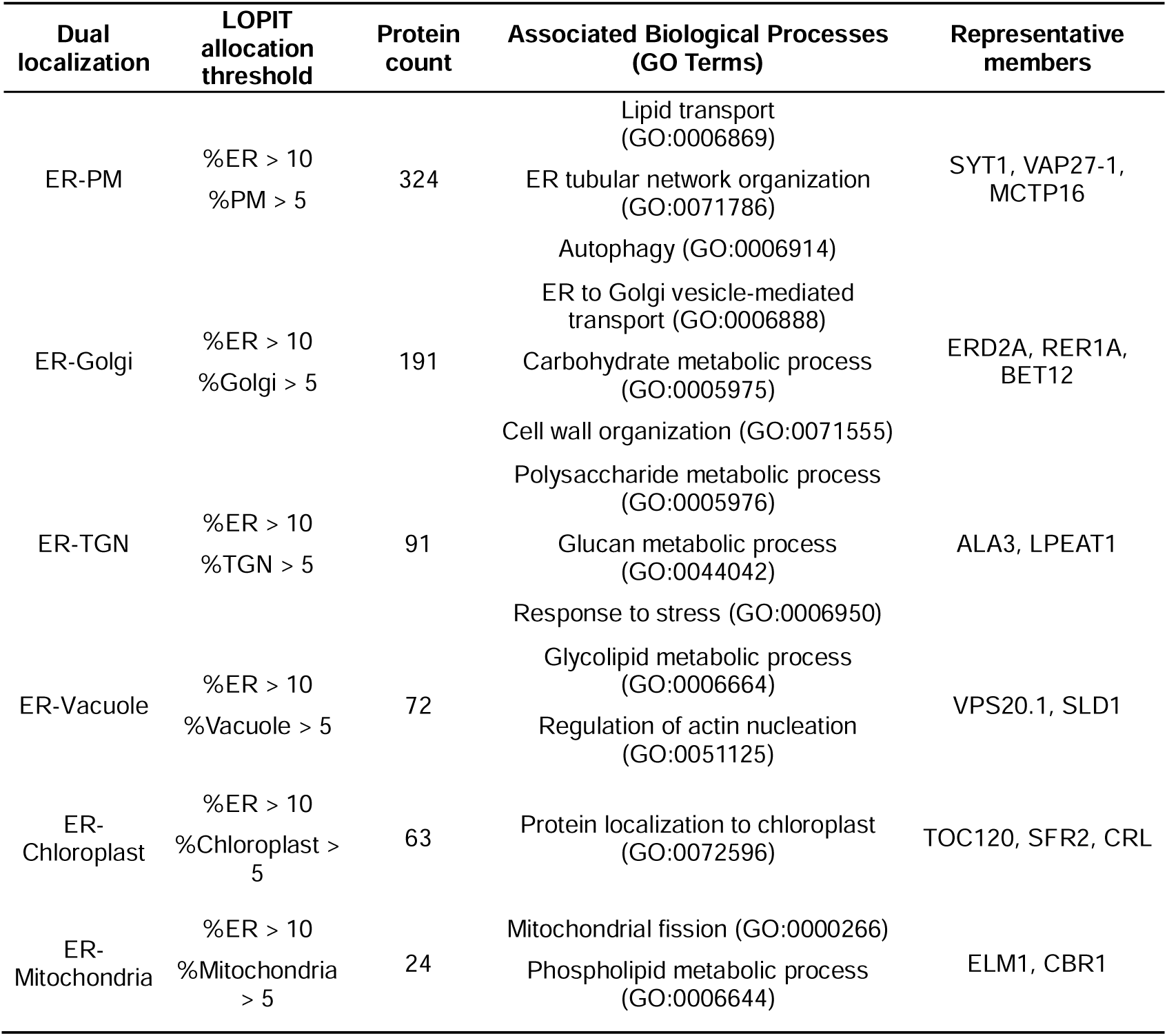
LOPIT dual-localization yields candidates and defines the functional signatures of diverse ER–organelle interfaces. Associated biological processes for each interface were determined through Gene Ontology (GO) enrichment analysis. Selected GO terms and representative members were chosen based on functional association with processes operating at each specific subcellular location.

Dual localization data revealed a clear functional specialization across the diverse contact boundaries (**Table 1**). Remarkably, lipid transport and metabolism^18,38,48,49^, cell wall biosynthesis^30,50^, and membrane trafficking^34,39^ are central within the operational landscape of these junctions. Within the ER-PM interface, enriched functional annotations largely mirror the profile of the SYT1 interactome, while also uncovering processes occurring at these interfaces that may not necessarily involve SYT1, such as autophagy^49,51^. Moving along the secretory pathway, the ER-Golgi and ER-TGN interfaces emerged as the next largest groups, possibly due to their structural similarity to the ER-PM interface^52^, cycling of trafficking vesicles^45^ and their role as a continuum for membrane remodeling and cell wall matrix deposition^52,53^. In contrast, the significantly lower protein density recovered at the ER-Mitochondria and ER-Chloroplast boundaries may indicate that interfaces with endosymbiotic organelles are fundamentally less dynamic. Because these endosymbiotic compartments do not participate in classical vesicular transport^52^, their junctional proteomes reflect a reduced presence of proteins that transiently flow, transit, or partition between these organelle membranes, pointing rather toward tighter or more static protein configurations.

### The SYT1 interactome correlates with LOPIT spatial proteomics and validates preferential ER-PM dual localization of interactors

To assess the spatial context of our AP-MS and PL interactomes, we cross-referenced the 289 identified SYT1 interactors with the HyperLOPIT spatial proteomics dataset^47^. We examined the distribution of the SYT1 interactome mapped onto a targeted organelle proteome subset containing the early secretory pathway (ER, PM, Golgi, and TGN), visualized via the DNN latent space analysis available from Choragraph^47^. The overlay of the protein distribution across these selected organelle clusters (**Fig. 2a**) with specific ER and PM localization (**Fig. 2b**) revealed a striking convergence, with the bait SYT1 mapping precisely to the interface bridging the ER and PM sectors. Mapping AP–MS and PL candidates onto this projection also showed that SYT1 interactors preferentially cluster within the ER and PM domains, supporting their spatial enrichment at the ER-PM contact sites (**Fig. 2c**). This spatial consistency demonstrates that the SYT1 interactome independently validates the proportional organelle assignments generated by Choragraph from the HyperLOPIT dataset.

**Figure 2.**
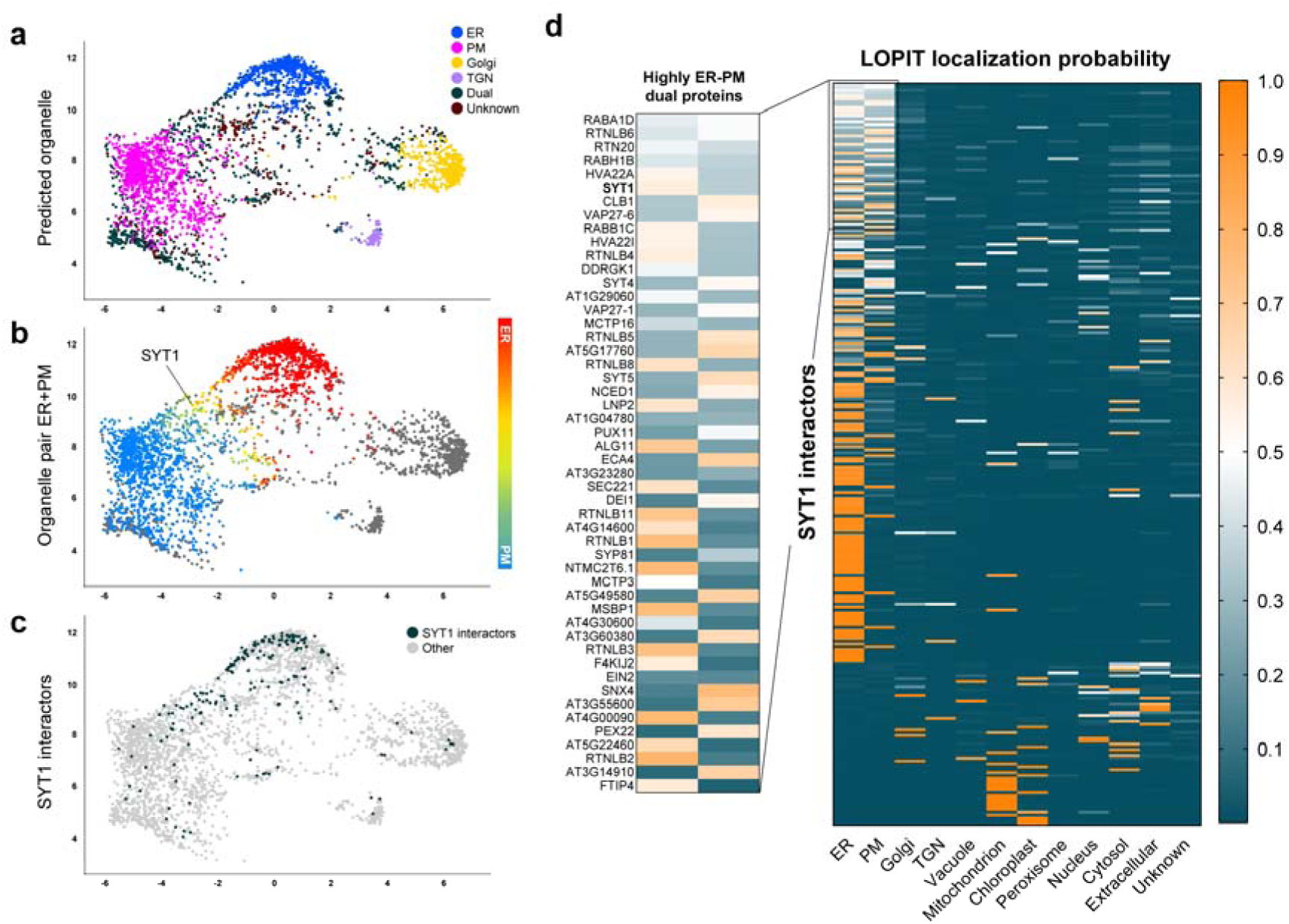
LOPIT dual localization proteomics reveals that SYT1 interactors are preferentially, but not exclusively, located at the ER and PM interfaces. **a, Spatial mapping of SYT1 interactors within a targeted organelle proteome.** Projection of a subset of the *Arabidopsis* spatial proteome (ER, PM, Golgi, TGN) derived from LOPIT data and visualized via Deep Neural Network (DNN) latent space analysis. Top panel: Distribution of all proteins across the selected organelle clusters. Middle panel: Visualization of ER and PM localization probabilities within the latent space; the color gradient highlights the confidence of proteins for these specific compartments. The SYT1 location within the projection is indicated. Bottom panel: Mapping of AP-MS and TurboID SYT1 interactors onto the targeted UMAP space. The clustering of SYT1-interacting proteins specifically within the ER and PM sectors illustrates a preferential enrichment at these subcellular domains. **b, LOPIT-based organelle localization probabilities of SYT1 interactors.** Heatmap showing the probability of localization of SYT1-interacting proteins to different organelles, as determined by LOPIT data. The color scale represents the probability of a given protein being localized to a specific organelle, ranging from 0 (absent, blue) to 1 (maximum probability, orange). Proteins are ordered according to an ER-PM Dual Localization Index, which prioritizes proteins with similar ER and PM localization probabilities and values close to 0.5. The inset highlights the subset of proteins with the highest ER-PM dual localization scores, which are candidates with the strongest predicted dual association with ER and PM.

Next, we employed these localization probabilities to categorize SYT1 interactors based on their HyperLOPIT specific membrane affinity, generating a localization heatmap of the SYT1 interactome (**Fig. 2d**). To identify candidates with higher probability to localize at ER-PM contact sites, proteins within this matrix were ordered according to an ER-PM Dual Localization Index, which prioritizes proteins with balanced, mid-range assignment values (close to 0.5; 50% composition) for both membranes. The detailed integration of the SYT1 interactomic and LOPIT localization datasets is provided in **Supplementary Data 3.** As structural tethers and core-resident components of ER-PM contact sites will exhibit similar affinities for the ER and the PM, applying this index enabled us to identify a cohort of high-scoring dual-affinity interactors (**Fig. 2d**). Remarkably, the highest-ranking candidates, which cluster at the top of the matrix, are strongly enriched for canonical tethering complexes and ER-shaping factors. For example, SYT1 itself emerged as the sixth most dual-localized protein in the entire list, directly validating the index ability to isolate *bona fide* ER-PM contact sites proteins.

### Multi-omics integration decodes uncharacterized machinery at ER-PM contact sites

To further investigate the molecular convergence between SYT1 interactomics and spatial proteomics, we compared the filtered AP-MS and PL datasets (enrichment score ≥ 20) with the 324 high-confidence dual ER-PM proteins extracted from Choragraph’s predictions (**Fig. 3a**). A total of 60 proteins (18%) of the dual ER-PM candidates identified in the LOPIT were present in the SYT1 interactome. This corresponds to nearly one-fifth of all dual-localized proteins identified with a single bait, providing robust validation of the data and suggesting that SYT1 plays a central role in the organization and function of the plant ER-PM contact sites.

**Figure 3.**
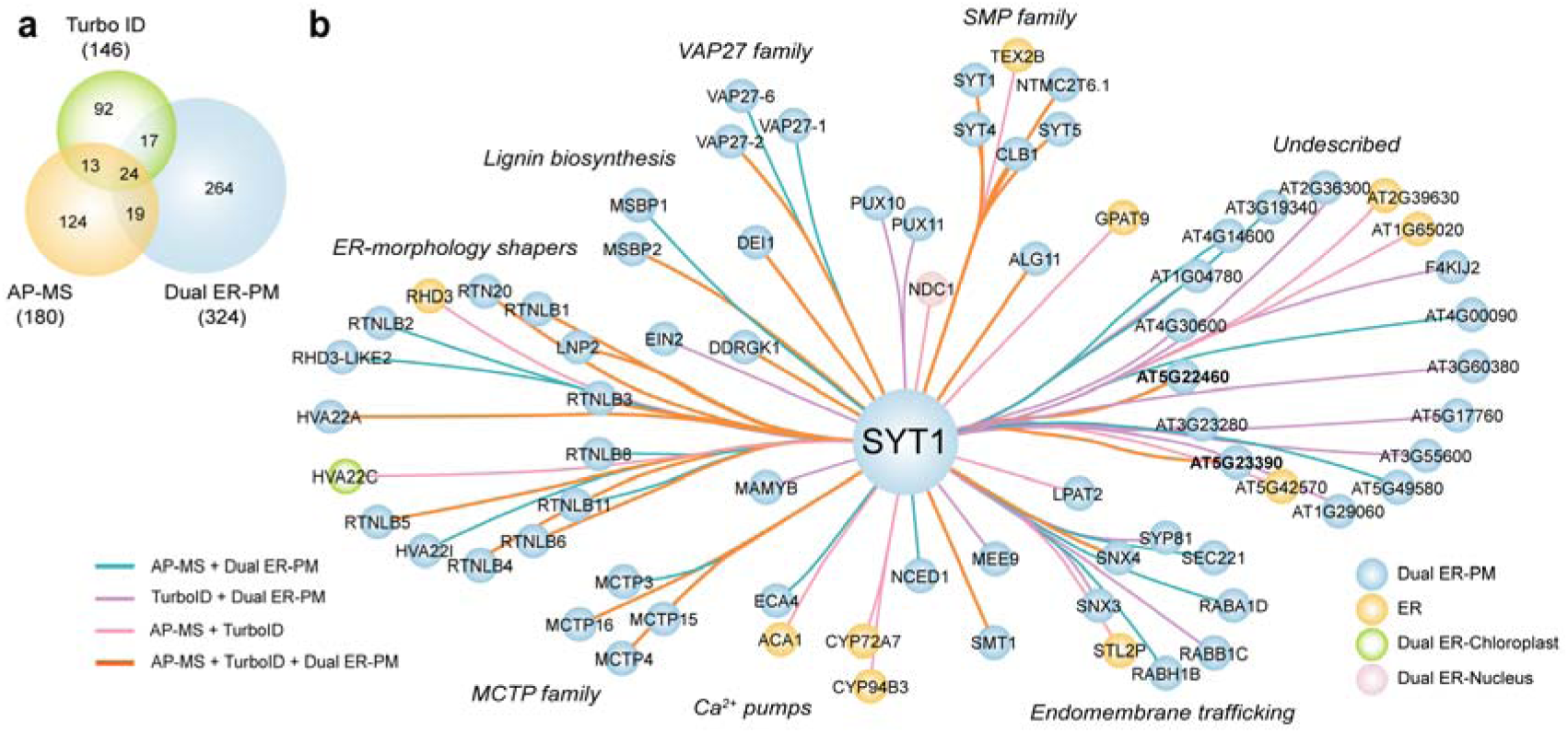
Integration of SYT1 interactomics and LOPIT dual-localization data defines the molecular landscape of ER-PM contact sites and uncovers uncharacterized candidates. **a, Intersection of the three proteomic datasets.** Venn diagram displaying the intersection between SYT1 AP-MS and TurboID datasets with the ER-PM dual-localized proteins defined by LOPIT data. The threshold criteria for the interactomic data (TurboID and AP-MS) is an enrichment ≥ 20, while dual ER-PM localization is established using localization probabilities ≥ 10% for the ER and ≥ 5% for the PM. **b, Network topology and functional clustering.** Interactor network mapping the candidate proteins identified at the triple intersection of the datasets. Edge colors represent different overlapping classifications, while node colors indicate LOPIT-predicted subcellular localizations. Proteins are organized into clusters following functional and structural criteria. Bolded entries within the "Undescribed" cluster indicate the uncharacterized candidates selected for downstream functional validation.

Next, we mapped the 60 shared candidates into an integrated interaction network (**Fig. 3b**). Beyond recovering the structural backbone of ER-PM contact sites (SMP, VAP27, and MCTP families, together with ER-morphology shapers), these new candidates could provide molecular insights into functional roles, e.g. Ca^2+^ homeostasis, that are well-characterized in animal MCS but remain largely unexplored in plants^54^. For instance, this analysis identified members from both major families of ER-anchored Ca^2+^ pumps: ACA1 and ECA4^55^ (**Fig. 3b**). Interestingly, data integration identified the membrane steroid-binding proteins MSBP1 and MSBP2, which are involved in scaffolding key enzymes of lignin biosynthesis^56^ (**Fig. 3b**). This suggests that SYT1 may function to anchor the phenylpropanoid biosynthetic machinery at the cell cortex, a link investigated later in the manuscript. Finally, the network revealed a substantial cluster of poorly characterized proteins. Among these are AT5G22460 and AT5G23390 (**Fig. 3b**, bolded), whose proteins were recovered across all three independent datasets and selected for further validation.

### Subcellular and biochemical characterization identifies SEPC1 and SEPC2 as *bona fide* ER-PM contact site components

To begin characterizing these novel ER-PM candidates, we performed *in silico* AlphaFold 3 structural predictions for AT5G22460 and AT5G23390 (**Fig. 4a**). Based on their robust physical and spatial association with SYT1, we named them **S**YT1-associated **E**R-**P**M **C**ontact Site proteins: SEPC1 (AT5G22460) and SEPC2 (AT5G23390). SEPC1 encodes a 340-amino-acid protein characterized by an N-terminal signal peptide followed by an α/β-hydrolase fold, while SEPC2 encodes a 730-amino-acid protein with several transmembrane domains. Neither sequence appears to contain lipid-binding modules (such as polybasic patches or C2 or PH domains)^5,41^, nor established protein-protein tethering motifs (such as MSP domains)^28^, which are typically required for autonomous recruitment to membrane junctions^4,14^.

**Figure 4.**
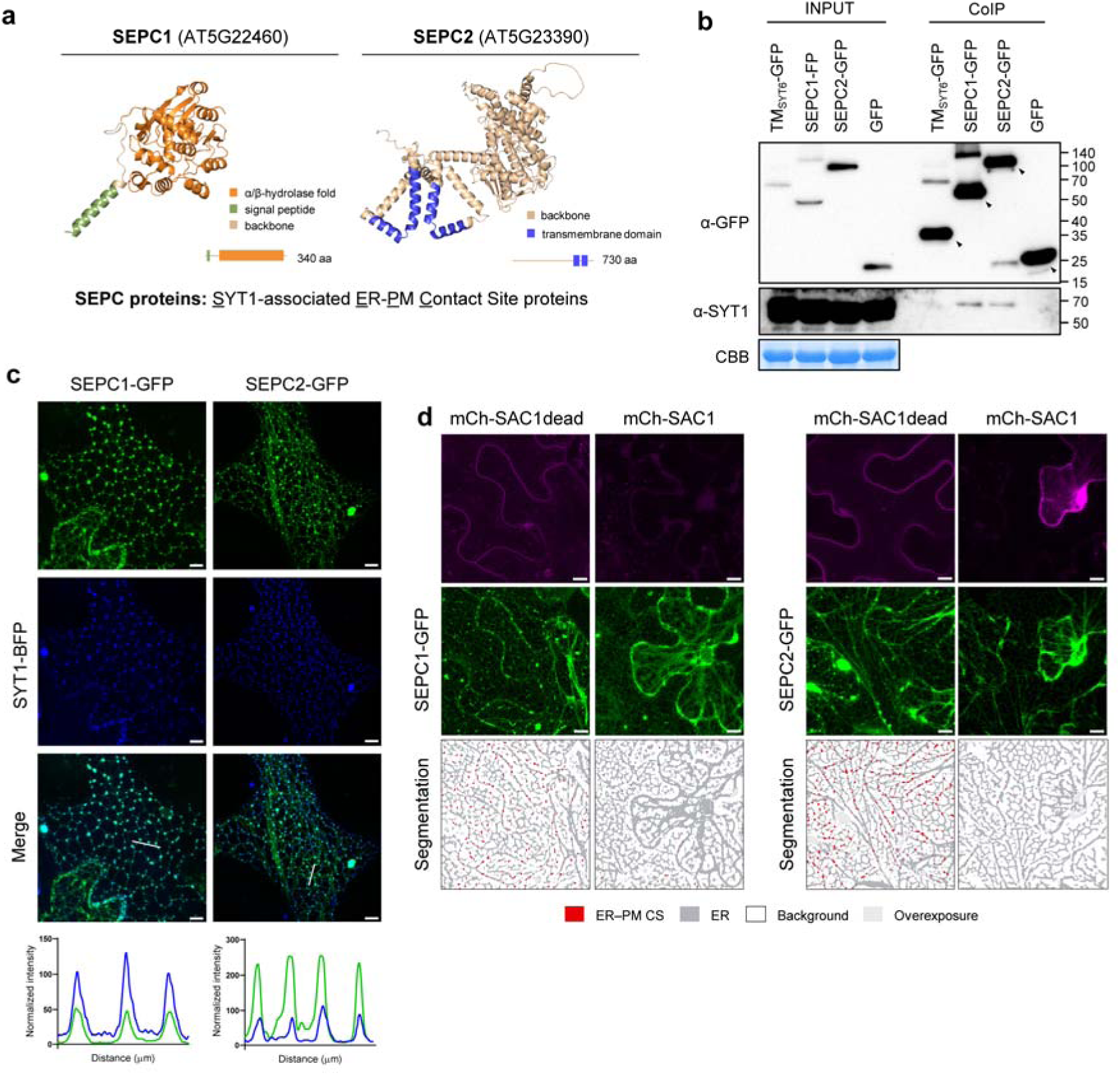
The SYT1-Associated Contact Site (SEPC) proteins are newly discovered components of ER-PM contact sites. **a, Structural characterization of novel SEPC proteins.** AlphaFold 3 structural predictions of uncharacterized candidates (SEPC1 and SEPC2). Diagrams illustrate predicted functional domains (orange), transmembrane segments (blue), and signal peptides (green). **b, Co-immunoprecipitation validation of SEPC proteins with SYT1.** *In vivo* validation of SYT1-SEPC1/2 association using transiently transfected *Arabidopsis* protoplasts. SEPC proteins were co-immunoprecipitated following GFP pull-down under non-denaturing conditions; bands corresponding to co-immunoprecipitated proteins are indicated by arrowheads. Soluble and ER-anchored GFP served as negative controls. Coomassie Brilliant Blue (CBB) staining of input samples served as a loading control. **c, Subcellular co-localization of SEPC proteins with SYT1.** Confocal micrographs displaying the cortical distribution of newly characterized SEPC proteins alongside SYT1 in transiently expressed *N. benthamiana* epidermal cells. Plot profile analysis (white line, slightly offset for clarity) of both candidates illustrates signal co-accumulation at SYT1-positive foci. Scale bars, 5 µm. **d, Dependence of SEPC localization on plasma membrane PI4P.** Confocal images of SEPC proteins following SAC1-driven depletion of plasma membrane PI4P, in transiently expressed in *N. benthamiana* epidermal cells. The loss of cortical foci under PI4P depletion confirms their classification as ER-PM contact site components. A catalytically inactive version of the enzyme (SAC1dead) served as a negative control treatment. Lower panels display Ilastik-based signal segmentation highlighting punctate (red) versus reticulated (gray) fluorescence patterns. Scale bars, 5 µm.

To further investigate the association between SYT1 and SEPC1/SEPC2, we performed co-immunoprecipitation (CoIP) assays using *Arabidopsis* mesophyll protoplasts transiently expressing SEPC1-GFP or SEPC2-GFP (**Fig. 4b**). This system preserves the native cellular environment, allowing the pull-down of endogenous SYT1 using a specific antibody. Furthermore, to rule out nonspecific associations due to ER membrane trapping, we used both a soluble GFP and an ER-targeted GFP (TM-SYT6-GFP) as controls. The latter consists of GFP fused to the transmembrane domain of SYT6, a plant synaptotagmin absent from the SYT1 interactome. CoIP assays confirmed that SEPC1 and SEPC2 pull down endogenous SYT1, but not the controls (**Fig. 4b**), demonstrating their *in vivo* association. Confocal imaging of SEPC1-GFP and SEPC2-GFP in protoplasts showed puncta of variable size that, especially in the case of SEPC2-GFP, resembled the morphology of ER-PM contact sites (**Supplementary Fig. 4**).

Next, we performed confocal microscopy to determine the subcellular localization of SEPC1 and SEPC2 *in planta*. When ectopically expressed in *N. benthamiana*, both SEPC1-GFP and SEPC2-GFP displayed a punctate cortical distribution characteristic of ER-PM contact sites, which perfectly overlapped with SYT1-BFP-marked foci (**Fig. 4c**). To further investigate the molecular basis of this localization, we tested the importance of phosphoinositides for protein targeting, focusing on phosphatidylinositol-4-phosphate (PI4P). This lipid is crucial for the PM recruitment of ER-PM contact site proteins, including SYT1 (see companion manuscript, Marković et al., 2026). To determine whether SEPC PM localization is dependent on PI4P, we transiently co-expressed SEPC1 and SEPC2 proteins with a PM-anchored version of the yeast PI4P phosphatase MAP-mCh-SAC1 (SAC1), using the catalytically inactive MAP-mCh-SAC1dead (SAC1dead) as a control^57^ (**Fig. 4d**). Expression of active SAC1, but not the catalytically inactive version SAC1dead, promoted the depletion of this lipid at the PM and the relocalization of SYT1 from ER-PM contact sites to the bulk ER^18,57^ (see companion manuscript, Marković et al., 2026). Consistent with this, the expression of active SAC1 resulted in a severe reduction of cortical puncta of both SEPC1 and SEPC2 and their relocalization to the bulk ER (**Fig. 4d**). In contrast, the expression of SAC1dead did not alter the cortical distribution of either SEPC protein. Altogether, these results validate SEPC1 and SEPC2 as genuine residents of plant ER-PM contact sites, demonstrating that their cortical recruitment relies, either directly or indirectly, on the same plasma membrane lipid identity required for SYT1 tethering.

### SYT1 mediates the recruitment of the phenylpropanoid biosynthetic machinery to ER-PM contact sites

Next, we sought to identify novel biological functions of SYT1 using our dataset. Interestingly, we found an enrichment of proteins responsible for the phenylpropanoid pathway driving the biosynthesis of monolignol precursors required for lignin deposition (**Fig. 5a-c**). Monolignol biosynthesis involves the three core ER-resident cytochrome P450 monooxygenases: cinnamate 4-hydroxylase (C4H), p-coumarate 3’-hydroxylase (C3’H), and ferulate 5-hydroxylase (F5H)^56,58^ (**Fig. 5a**, **Supplementary Fig. 5**). These three enzymes are physically linked into a metabolon by Membrane Steroid Binding Proteins 1 and 2 (MSBP1 and MSBP2), which act as a structural scaffold to optimize metabolic flux^56^. Our interactomic datasets revealed that SYT1 physically associates with this structural scaffold across both TurboID PL and AP-MS assays (**Fig. 5b,c**). Furthermore, both MSBP1 and MSBP2 exhibit significant dual ER-PM localization probabilities within the HyperLOPIT spatial proteomics (**Fig. 5c**). The core cytochrome P450 enzymes C4H and C3’H were also highly enriched specifically within the AP-MS dataset, confirming a robust link with the SYT1 complex (**Fig. 5b, c**). In contrast, the downstream enzyme F5H was absent from our SYT1 interactomics and LOPIT datasets. This was somewhat expected as F5H generates syringyl (S) precursors in lignifying tissues^59,60^, and is unlikely to be expressed in undifferentiated cell cultures lacking secondary cell wall^61^.

**Figure 5.**
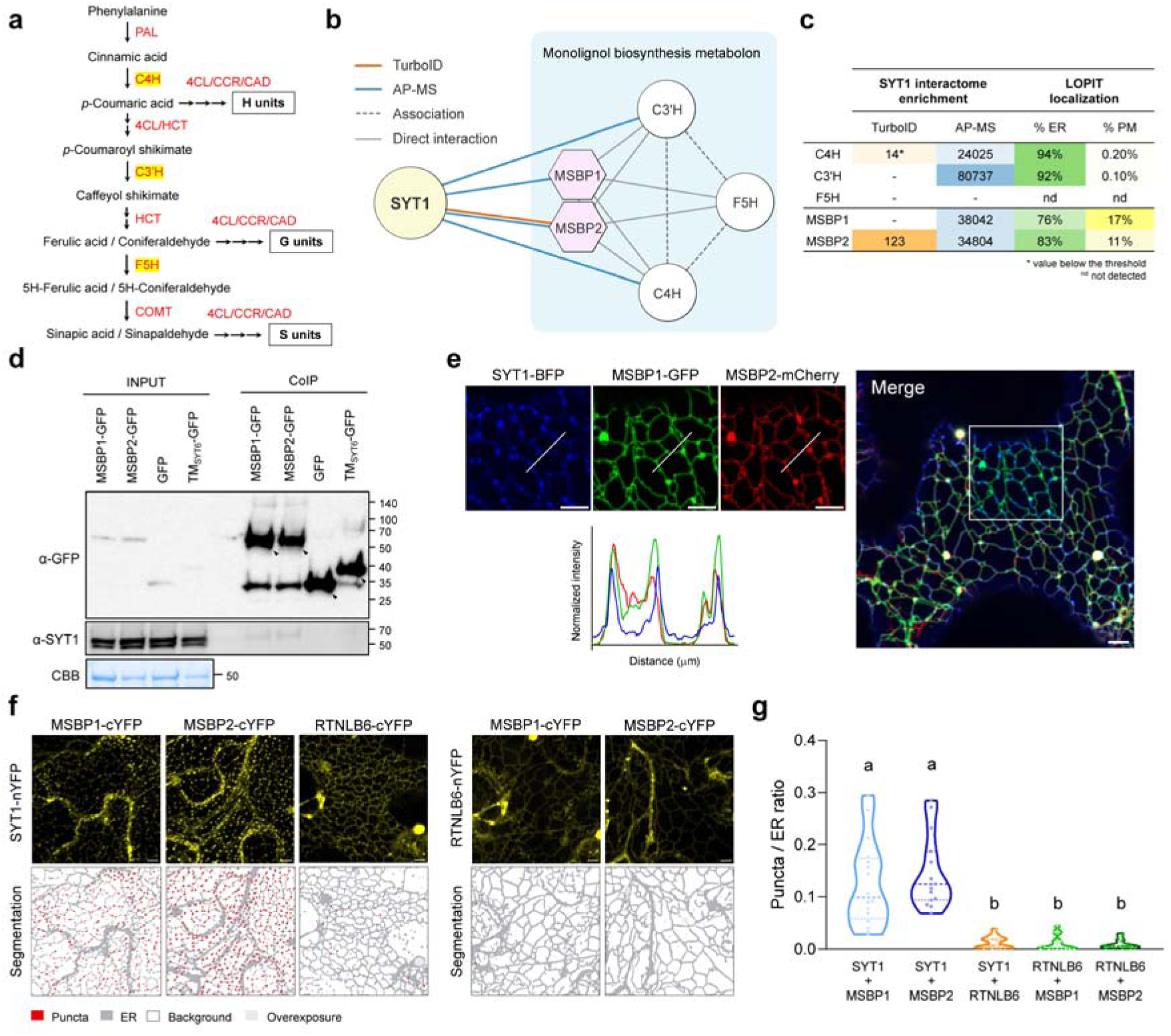
SYT1 anchors and organizes the 1 enzymatic cluster at ER-PM contact sites. **a, Schematic representation of the monolignol biosynthesis pathway.** Simplified diagram illustrating the position of P450 enzymes (C4H, C3’H, and F5H) within the phenylpropanoid pathway leading to lignin precursors. A complete diagram of the pathway can be found in **Supplementary** Figure 5. Cytochrome P450 monooxygenases investigated in this study (C4H, C3’H, and F5H) are highlighted in yellow boxes. Abbreviations: PAL, phenylalanine ammonia-lyase; C4H, cinnamate 4-hydroxylase; 4CL, 4-coumarate-CoA ligase; CCR, cinnamoyl-CoA reductase; CAD, cinnamyl alcohol dehydrogenase; HCT, hydroxycinnamoyl-CoA:shikimate hydroxycinnamoyl transferase; C3’H, p-coumarate 3’-hydroxylase; F5H, ferulate 5-hydroxylase; COMT, caffeic acid O-methyltransferase; H, p-hydroxyphenyl unit; G, guaiacyl unit; S, syringyl unit. **b, Protein-protein interaction network of SYT1 and the monolignol biosynthesis protein complex.** Nodes represent proteins and edges represent interactions. Colored edges denote experimental data (orange, TurboID; blue, AP-MS); gray edges indicate previously reported literature interactions (solid lines, direct interaction; dashed lines, association). Hexagonal nodes represent the homo- and heterodimerization of MSBP1/MSBP2. **c, Quantitative SYT1 interactome data and localization probabilities.** Summary table showing enrichment values obtained in TurboID and AP-MS assays from proteins within the complex. Subcellular ER and Plasma Membrane localization probabilities are based on LOPIT data. The asterisk (*) denotes detections below the established enrichment threshold. **d, Co-immunoprecipitation of SYT1 with MSBP1 and MSBP2.** *Arabidopsis* protoplasts were transiently transfected with MSBP1-GFP or MSBP2-GFP, and endogenous SYT1 was detected using specific antibodies following GFP pull-down under non-denaturing conditions. Soluble and ER-anchored GFP served as negative controls. Coomassie Brilliant Blue (CBB) staining of input samples indicates equal loading. **e, Subcellular co-localization of SYT1 with MSBP1 and MSBP2.** Confocal images displaying the subcellular distribution of SYT1, MSBP1, and MSBP2 transiently co-expressed in *N. benthamiana* leaves. While MSBPs are broadly distributed throughout the ER network, plot profile analysis illustrates signal co-accumulation at SYT1-positive foci. The white line indicates the region analyzed in the plot profile (slightly offset for clarity). Scale bar, 5 µm. **f, Spatial restriction of SYT1-MSBP1/2 interaction at ER-PM contact sites.** Confocal images of Bimolecular Fluorescence Complementation (BiFC) assays between SYT1 and MSBP1/MSBP2 in transiently expressed *N. benthamiana* epidermal cells. SYT1-RTNLB6 and RTNLB6-MSBP1/MSBP2 interactions serve as a control for cortical and bulk ER distribution, respectively. Lower panels display Ilastik-based signal segmentation highlighting punctate (red) versus reticulated (gray) fluorescence patterns. Scale bars, 5 µm. **g, Quantification of BiFC fluorescence morphology.** Distribution of punctate versus reticulated signal patterns for the indicated BiFC pairs. Data are presented as violin plots with individual data points; horizontal lines indicate the mean and quartiles (*n* ≥ 15 images). Statistical significance was determined by a Brown-Forsythe ANOVA followed by Games-Howell *post-hoc* test for multiple comparisons (*P* < 0.001).

These findings led us to further investigate whether SYT1 could serve as a localized docking platform, thereby organizing the MSBP-scaffolded machinery at ER-PM contact sites. To test this, we first analyzed the association between SYT1 and the MSBP scaffolds *in vivo* by performing CoIP assays in *Arabidopsis* protoplasts (**Fig. 5d**), which confirmed that MSBP structural scaffolds co-immunoprecipitated endogenous SYT1 but not GFP nor TM-SYT6-GFP controls (**Fig. 5d**).

In *Arabidopsis* protoplasts, MSBP1-GFP and MSBP2-GFP displayed a reticulated pattern with irregularly shaped granules as previously reported^56^ (**Supplementary Fig. 6a**). To determine their subcellular localization and association with SYT1 *in planta*, we performed transient co-expressions in *N. benthamiana*. Strikingly, MSBP1 and MSBP2 exhibited selective enrichment at discrete, SYT1-positive cortical foci at ER three-way junctions (**Fig. 5e**), while showing a bulk ER distribution when expressed individually (**Supplementary Fig. 6b**) as already characterized^56^. In addition, these co-localized foci remained immobile over time (**Supplementary Video 1**), consistent with their presence at ER-PM contact sites^26,42^. To further determine the precise localization of SYT1-MSBP interaction, we used Bimolecular Fluorescence Complementation (BiFC) assays (**Fig. 5f**). Reconstitution of the YFP fluorescence signal revealed a distinct, mostly punctate pattern at the cell cortex for both SYT1-MSBP1 and SYT1-MSBP2 interaction pairs (**Fig. 5f**). To verify the spatial specificity of this association, we employed the ER protein RTNLB6 as a control for the localization of this interaction. The SYT1-RTNLB6 pair reconstituted a cortical ER network pattern, displaying significantly fewer puncta than the MSBP pairs (**Fig. 5f**), and showing that not all SYT1 interactions are restricted to ER-PM contact sites. Additionally, the RTNLB6-MSBP1 and RTNLB6-MSBP2 pairs yielded a faint, exclusively reticulated distribution without any ER-PM contact sites enrichment (**Fig. 5f**). Quantification of BiFC images using the machine-learning based software Ilastik^62^ further verified the robust enrichment of SYT1-MSBP complexes within punctate domains (**Fig. 5g**). Altogether, these data support that SYT1 is part of the phenylpropanoid biosynthetic machinery at ER-PM contact sites through MSBP anchoring.

### SYT1 modulates ectopic lignification during cell wall integrity stress

To investigate if SYT1-mediated metabolic scaffolding of the monolignol biosynthesis machinery is physiologically relevant, we examined SYT1 behavior during conditions that induce ectopic lignification. To this end, we treated 5-days-old seedlings of a complementing *pSYT1:SYT1-GFP Arabidopsis thaliana* line in *syt1* background (hereafter SYT1-GFP) with the cellulose biosynthesis inhibitor isoxaben (600 µM, 18 h), a cell wall stress treatment that causes compensatory ectopic lignification in *Arabidopsis* roots^63,64^. Under mock conditions, root cells presented a regular shape with SYT1-GFP accumulated in the cortical region in a rather diffuse pattern (**Fig. 6a**). In contrast, isoxaben treatment triggered radial cell swelling, extensive accumulation of lignin in the apoplast and more punctate SYT1-GFP signal across the entire PM interface, matching the onset of lignin deposition (**Fig. 6a,b**). This indicates that cell wall stress caused by isoxaben drives a dynamic reorganization of SYT1 into ER-PM contact sites in coordination with active lignification.

**Figure 6.**
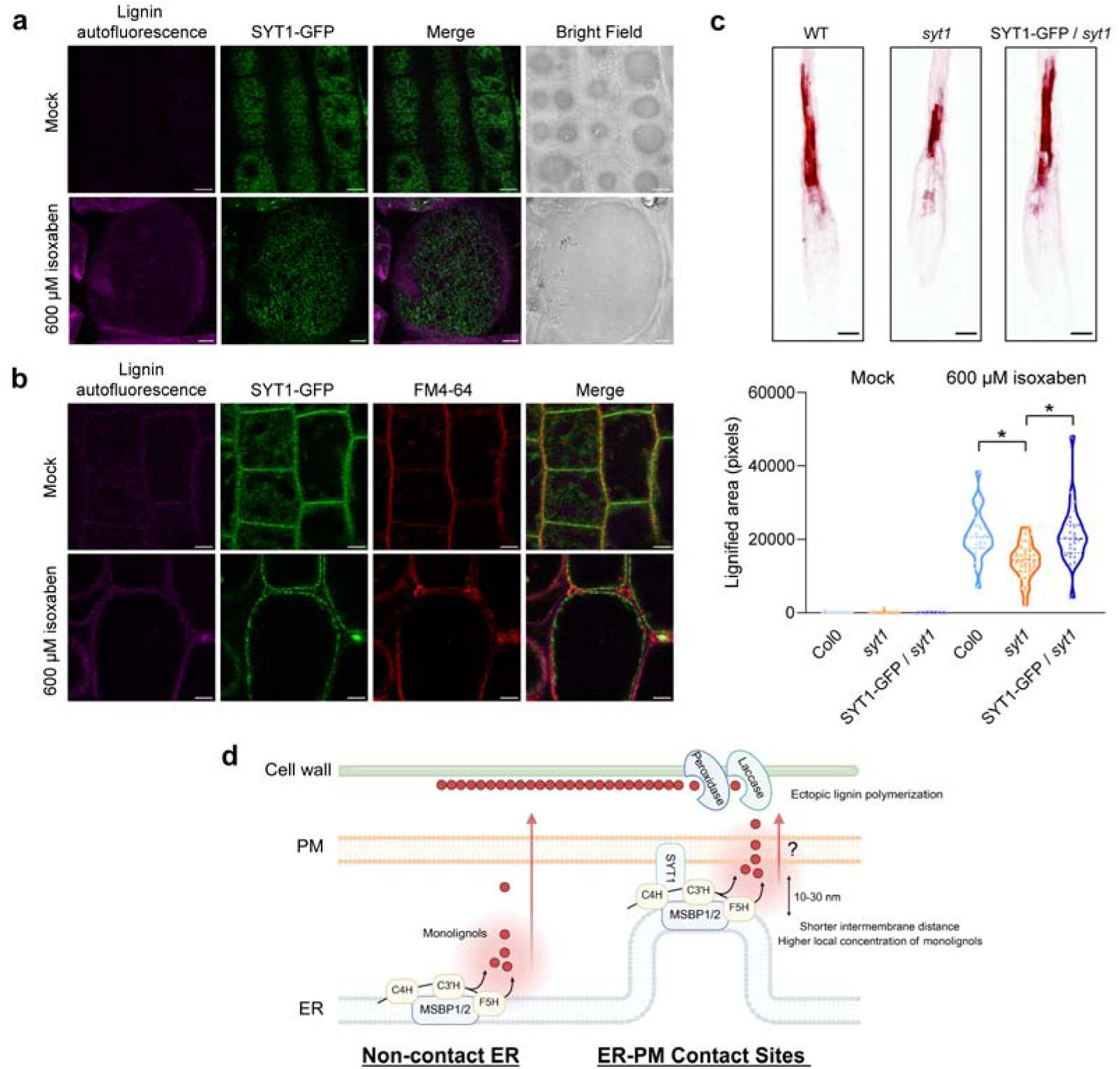
SYT1 is required for ectopic lignification during cell wall integrity stress. **a, Cortical plane visualization of SYT1 contact site remodeling and cell hypertrophy under cell wall stress.** Confocal images displaying a surface cortical view of Arabidopsis root cells expressing SYT1-GFP under mock control (top) or isoxaben-treated (600 µM, 18 h; bottom) conditions. Ectopic lignin deposition was tracked via autofluorescence (magenta; Exc. 405 nm / Em. 510–580 nm). Transmitted light (Bright field) channel is included to illustrate radial cell swelling. Scale bars, 5 µm. **b, Isoxaben-induced SYT1 relocalization on ectopically lignified root cells.** Confocal images of root cells expressing SYT1-GFP under mock (top) and isoxaben-treated (600 µM, 18 h; bottom) conditions. SYT1-GFP transitions from a diffuse ER pattern to a punctate ER-PM contact sites pattern, which correlates with the spatial onset of ectopic lignin deposition. Lignin was visualized via autofluorescence (magenta; Exc. 405 nm / Em. 510–580 nm) and membranes were stained with FM4-64 (red). Scale bars, 5 µm. **c, Defective ectopic lignification in *syt1* mutant following isoxaben treatment.** (Top) Representative images of phloroglucinol-HCl staining on roots of WT, *syt1*, and SYT1-GFP seedlings subjected to cell wall stress (600 µM isoxaben, 18 h). Scale bars, 100 µm. (Bottom) Quantitative machine-learning analysis of lignin deposition in Col0, *syt1*, and *pSYT1:SYT1-GFP* complemented seedlings. Loss of SYT1 results in a significant reduction in stress-induced lignification compared to wild-type and complemented lines. Data are presented as violin plots with individual data points; horizontal lines indicate the mean and quartiles (treatment: *n* ≥ 25 images; mock: *n* ≥ 10 images). Statistical analysis was performed among genotypes within each group. Statistical significance was determined by a Brown-Forsythe ANOVA followed by a Games-Howell post-hoc test for multiple comparisons. Single asterisks (*) denote pairs with significant differences (*P* < 0.001). **d, Working model for SYT1-mediated metabolon anchoring during cell wall stress.** Schematic representation of the proposed mechanism for SYT1-mediated tethering of the MSBP1/2 metabolon at ER-PM contact sites. SYT1 acts as a molecular tether that recruits the MSBP-scaffolded P450 metabolon (comprising C4H, C3’H, and F5H) to ER-PM contact sites. This organization narrows the intermembrane distance (10-30 nm), facilitating a high local concentration of monolignols for efficient export and subsequent polymerization into the cell wall by laccases and peroxidases.

Finally, to assess whether SYT1 plays a role in lignin deposition under isoxaben-triggered stress, we evaluated ectopic lignification in the wild-type (WT), *syt1* mutant, and *pSYT1:SYT1-GFP* complementation lines using phloroglucinol-HCl staining^65^. This analysis revealed a reduced lignified area in the roots of *syt1* mutants compared to WT and complementing seedlings after an 18-h treatment with 600 µM isoxaben (**Fig. 6c**, **top**). These differences were quantified using automated machine-learning-based signal segmentation to detect phloroglucinol-stained regions in an unbiased manner (**Supplementary Fig. 7**). Quantitative analysis indeed confirmed a statistically significant reduction in the total lignified root area of *syt1* after isoxaben treatment compared to WT and the complementing line (**Fig. 6c**, **bottom**). Moreover, dose-response assays revealed that the defective lignification of *syt1* was less pronounced at lower isoxaben concentrations (**Supplementary Fig. 8**). Thus, these results indicate that SYT1 is required to sustain efficient lignin deposition when monolignol biosynthetic demand increases.

## DISCUSSION

In this work, we studied the interaction landscape of the ER-PM contact sites tether SYT1 to uncover cellular processes associated with these nanodomains in plants. The use of both AP-MS and TurboID provided complementary information on proteins that either associate with SYT1 or are localized at the ER-PM contact sites due to their proximity to SYT1. The recovery of known ER-PM contact site components, including members of VAP27^27,28,66,67^, MCTP^14,30^, and SYT^17,18,42^ families, supports the reliability of our dataset. At the same time, the identification of proteins not previously linked to MCS indicates that the molecular composition of plant ER-PM contact sites is far from complete (**Fig 1 a,b**). The enrichment of proteins associated with ER organization, vesicle trafficking, calcium transport, lipid binding, sterol metabolism, and heme-containing monooxygenase activity suggests that SYT1-associated contact sites host a diverse set of cellular activities beyond membrane tethering or lipid transport.

The integration of interaction datasets with LOPIT spatial proteomics enabled us to further refine the SYT1 interactomic landscape (**Fig. 3**), while the ER-PM dual localization index provides a useful prioritization tool. Sixty SYT1-associated proteins displayed localization compatible with dual ER and PM association. Among them are several RTNLBs that shape ER morphology previously identified as SYT1 interactors^35^. Given that SYT1 preferentially localizes at highly curved cortical ER regions^17,26^, interactions with RTNLBs may help position SYT1 at ER tubules or cisternal edges, allowing its C2 domains to contact the PM and stabilize these nanodomains. SYT1 also interacted with other ER morphology regulators, including LNP1 and LNP2^68,69^, RHD3, and RHD3-LIKE2^37^, consistent with a role for SYT1 in ER organization^13,26^. Importantly, a localization outside the strict ER-PM interface does not exclude functions at contact sites. Examples include DGK1 and DGK2 which, despite showing bulk ER localization in *N. benthamiana*, interact with SYT1 specifically at the ER-PM contact sites^18^. Indeed, the association with ER-PM contact sites can happen in transient manner, as demonstrated with the dynamic association of PI4P phosphatase SAC7 with SYT1 (see companion manuscript Marković et al., 2026). In addition, proteins localized at the ER-PM contact sites may have additional localizations and functions. For example, VAP27 proteins, considered typical ER-PM contact sites tethers, have recently been identified at ER-chloroplast MCSs, forming a functional complex with ORP2A to regulate sterol levels in chloroplasts^67^, and also at ER-lipid droplet contact sites, regulating their biogenesis^70^.

Notably, the identification of ECA1 and ACA1 as SYT1 interactors is particularly interesting, as they are ER-localized Ca^2+^ ATPases that may contribute to Ca^2+^ homeostasis at ER-PM contact sites^71,72^. In mammalian cells, the SERCA Ca^2+^ ATPases have been linked to ER-PM contact sites during store-operated Ca^2+^ entry, where they colocalize with STIM1/Orai1 puncta and promote efficient refilling of ER Ca^2+^ stores^54^. Although the role of plant ER-PM contact sites in Ca^2+^ signaling and regulation has long been postulated, little evidence has been provided about Ca^2+^ transporters that may function at these nanodomains in plants^73^. The architectural clustering of these plant Ca^2+^ pumps could represent a direct mechanism linking ER-PM contact sites with the intracellular regulation of Ca^2+^ signaling. Thus, this compartmentalized control may point to the long-sought connection between plant ER-PM contact sites and cellular Ca^2+^ signaling, specifically during stress conditions that trigger robust intra- and intercellular Ca^2+^ waves^74,75^. In addition, given the calcium-dependent lipid-binding nature of Synaptotagmin C2 domains^41^, it is possible that local Ca^2+^ transport might in turn feed-back on ER-PM contact site organization.

As proof of principle, we selected two uncharacterized candidates from this high-confidence group, SEPC1 and SEPC2, for experimental validation. Their association with SYT1 was confirmed by co-immunoprecipitation (**Fig. 4b**), and both proteins accumulated in cortical puncta that overlapped with SYT1-positive foci (**Fig. 4c**). Moreover, PM depletion of PI4P by SAC1 disrupted their punctate localization, confirming that their mechanism for PM targeting is comparative to the ER-PM contact site proteins (**Fig. 4c**). These results expand the known molecular repertoire of plant ER-PM contact sites and indicate that other uncharacterized proteins within the SYT1 protein network may contribute to contact-site architecture, regulation, or specialization.

ER-PM contact sites have been primarily associated with lipid transfer^17,18^ and stress responses^19,20,33,42^. An interesting finding was the association between SYT1 and proteins involved in monolignol biosynthesis, a plant-specific process (**Fig. 5a,b**). Notably, this connection had already been hinted at in a previous AP-MS proteomic study that used the cytochrome P450 enzymes of the phenylpropanoid pathway, C4H and C3’H, as baits^76^ (**Supplementary Table 2**). They identified SYT1, VAP27, and other proteins also present in our proteome, providing another proof of the link between ER-PM contact sites and lignin precursors metabolism.

To clarify how this biosynthetic machinery is organized at the ER-PM interface, we focused on MSBP1 and MSBP2, as they are established interactors of this core cytochrome P450 enzymes^56^ and emerged as prominent hits in our SYT1 spatial dataset. Confocal microscopy showed local co-accumulation of these proteins at discrete SYT1-positive cortical foci (**Fig. 5e**, **Supplementary Video 1**). BiFC assays indicated that SYT1-MSBP interactions are spatially restricted to ER-PM contact sites, in contrast to the more reticulated ER pattern observed for RTNLB6-MSBP interactions (**Fig. 5f,g**). This difference suggests that MSBP proteins may participate in distinct ER-associated complexes, while SYT1 specifically recruits or stabilizes a subpopulation of MSBP1/2. Cell wall stress induced by isoxaben caused a redistribution of SYT1-GFP from a more diffuse ER pattern to punctate cortical structures associated with lignin autofluorescence, indicating dynamic remodeling of SYT1-positive ER-PM contact sites upon impaired cellulose biosynthesis (**Fig. 6a**). Moreover, *syt1* showed reduced ectopic lignification after isoxaben treatment, whereas SYT1-GFP restored this phenotype, indicating that SYT1 is required for efficient stress-induced lignification (**Fig. 6b,c**).

The spatial organization of the monolignol biosynthetic machinery discovered here provides critical insights into a long-standing debate in plant biology. While the synthesis of monolignols at the cytosolic ER face and their subsequent polymerization in the apoplast are well characterized^77^, the intervening transport mechanism across the plasma membrane has remained elusive, with active ABC-type transporters^78^, passive bilayer diffusion^79,80^, and the less-probable vesicular trafficking^80^ representing the primary co-existing hypotheses (**Supplementary Fig. 9**). Passive diffusion models inherently rely on a polymerization-induced concentration gradient, wherein the rapid conversion of monolignols into lignin in the cell wall creates a continuous metabolic sink across the plasma membrane^79,80^.

Taking all together, we propose a working model wherein SYT1-associated ER-PM contact sites act as organizing platforms for stress-induced lignification, providing a mechanistic bridge that anchors the monolignol biosynthetic metabolon at the ER-PM interface (**Fig. 6d**). In this model, the intermembrane 10-30 nm gap defines a highly confined functional nanodomain that minimizes diffusion distance and limits substrate dispersion. This unique junctional architecture promotes the local accumulation of lignin precursors directly adjacent to the plasma membrane-cell wall interface, thereby maximizing the concentration gradient between the cytosol and the apoplast. This spatial confinement drives a rapid, efficient, and passive non-vesicular export pathway precisely under stress conditions that require high lignification rates, setting up the structural basis for highly localized cell wall remodeling.

In summary, this study expands the map of plant ER-PM contact site proteins and identifies SYT1 as an organizer of a stress-responsive contact-site domain linked to monolignol biosynthesis. Indeed, the connection between ER-PM contact sites and phenylpropanoid metabolism supports an important role of these nanodomains in coordinating cellular architecture with adaptive cell wall responses. Under this new light, SYT1-associated ER-PM contact sites emerge as actors that do more than physically tether membranes or transport lipids; rather, they constitute central platforms where metabolic machinery is orchestrated for crucial roles such as cell wall reinforcement.

## MATERIALS AND METHODS

### Plant Material and Growth Conditions

*Arabidopsis thaliana* suspension cell cultures (Ler ecotype), *Arabidopsis* plants (Col-0 ecotype) and *Nicotiana benthamiana* plants were used as plant material for this study. *Arabidopsis* plant genotypes utilized in this work included wild type (Col-0 ecotype), *syt1* mutant (*syt1-2*, SAIL_775_A08) and complemented *pSYT1:SYT1-GFP* transgenic line, characterized in previous works^17,20^. For AP-MS and TurboID PL assays, *Arabidopsis* cell cultures were maintained under a long day-conditions regime at 22 °C on a shaker at 150 rpm, as previously described^81^. For seedlings germination, *Arabidopsis* seeds were surfaced-sterilized using chlorine vapors for 4 h and plated under sterility on half-strength Murashige-Skoog containing 1.5% (w/v) sucrose and 0.8% (w/v) agar. Seeded plates were vernalized for 2 days at 4°C in darkness and vertically grown in long-day conditions (16 h light/8 h dark, 130 ± 30 µmol photons m^−2^ s^−1^, 22±1 °C). Arabidopsis Col-0 plants for protoplast isolation were transferred to soil (organic substrate:vermiculite, 4:1 v/v) and grown in short-day conditions (8 light/16 h dark, 130 ± 30 µmol photons m^−2^ s^−1^, 22±1 °C).

### Molecular cloning

To amplify genomic and cDNA sequences, *Arabidopsis* Col-0 genomic and cDNA were used as templates for sequence amplification with the high-fidelity polymerase iProof (Bio-Rad #1725301). Genomic sequences of SEPC1 (AT5G22460), SEPC2 (AT5G23390), and MSBP1 (AT5G52240), alongside the coding sequence (CDS) of MSBP2 (AT3G48890), were cloned into entry vectors using the MultiSite Gateway system. Primers used for cloning are listed in **Supplementary Data 4**. All constructs were verified using colony PCR, restriction analysis, and sequencing.

Cloning procedures for vectors used in AP-MS and TurboID proximity labeling are detailed in their respective sections. For transient expression in *Nicotiana benthamiana*, constructs were assembled into the MultiSite Gateway destination vector FastRed pLOK180_pFR7m3 by recombining entry clones containing the UBQ10 promoter, the respective genomic or CDS sequences, and fluorescent tags (BFP, GFP, or mCherry). For expression in Arabidopsis mesophyll protoplasts, entry clones were recombined into a Gateway-adapted pUC19 destination vector containing the cauliflower mosaic virus (CaMV) 35S promoter and a GFP tag. For Bimolecular Fluorescence Complementation (BiFC) assays, sequences were cloned into the pDEST-GW-cYFP destination vector^82^ to generate the MSBP1-cYFP and MSBP2-cYFP constructs. Previously generated BiFC plasmids available in our laboratory, including SYT1-nYFP, RTNB6-nYFP, and RTNB6-cYFP^18^, were also employed in this work. All used constructs are detailed in **Supplementary Data 4**.

### Affinity Purification Coupled to Mass Spectrometry (AP-MS)

AP-MS experiments were made in the Interactomics Facility of the VIB department of Plant Systems Biology. The open reading frame (ORF) encoding the SYT1 protein (AT2G20990) was cloned into the pDONR221 vector, sequence-verified, and recombined with gateway entry vectors encoding CaMV 35S promoter and the C-terminal GS^rhino^ tag through multisite Gateway LR reaction into the pKCTAP destination vector. The resulting expression vector was transformed to *Agrobacterium tumefaciens* C58C1 (pMP90). *Arabidopsis thaliana* PSB-L cell suspension cultures were transformed with the constructs, and expression of SYT1-GS^rhino^ and membrane solubilization efficiency were initially evaluated in 50 mL pilot cultures by extraction with an isolation buffer (50 mM HEPES pH 7.5, 15 mM MgCl_2_, 150 mM NaCl, 15 mM p-nitrophenyl phosphate, 60 mM β-glycerophosphate, 0.1 mM Na_3_VO_4_, 1 mM NaF, 5% (v/v) Ethylene Glycol, 1 mM PMSF, 1 µM E64, cOmplete™ ULTRA EDTA-free Protease Inhibitor Cocktail (Roche)), supplemented with 1% (w/v) digitonin. Upon verification of robust and sufficient bait expression, cell cultures were scaled up to 2 L and harvested. For AP-MS, total membrane proteins were extracted using 1% (w/v) digitonin in the extraction buffer. Prior to affinity purification, extracts were treated with 3 mM dithiobis(succinimidyl propionate) (DSP) to stabilize proximal protein interactions via chemical crosslinking. After 45 min crosslinking on ice on a orbital shaker, non-reacted DSP was neutralized by addition of 1 mL 1 M Tris-HCl buffer (pH 7.5). Subsequently, three independent replicates of affinity purification (pull-down) were performed, as previously described with minor adaptations^31^. In brief, 25 mg total proteins were incubated for 2 h with 50 µL magnetic IgG beads. Next, beads were subsequently washed with 500 µL extraction buffer, 2x with 500 µL extraction buffer with 0.2% digitonin, and once with 500 µL extraction buffer without detergent. Proteins were eluted with 3x 150 µL 0.2 M Glycine (pH 2.5), reduced for 30 min with 5 mM TCEP at 37 °C and alkylated for 30 min with 10 mM Iodoacetamide. Finally, proteins were digested with Trypsin/Lys-C, and peptides were purified using C18 Omix ZipTips (Agilent), dried in a SpeedVac, and stored at −20 °C prior to MS analysis.

### Proximity Labeling (TurboID)

TurboID proximity labeling (PL) experiments were made in the Interactomics Facility of the VIB department of Plant Systems Biology. The open reading frame (ORF) encoding the SYT1 protein (AT2G20990) was cloned into the GreenGate pGGC000 vector, sequence-verified, and recombined with a C-terminally TurboID-tag and the CaMV 35S promoter through Gibson Assembly into pGGK-AG. Transgenic *Arabidopsis thaliana* PSB-D cell suspension lines were generated, and SYT1-TurboID production was confirmed in 50 mL pilot cultures. Following validation of proper bait accumulation, cultures were scaled up to 1 L. For *in vivo* proximity labeling, the cultures were treated with 50 μM biotin for 1 h at 25 °C immediately prior to harvesting. Streptavidin-based affinity purification of biotinylated proteins was performed in triplicate following an optimized TurboID purification protocol as previously described^32^.

### LC-MS/MS

For AP-MS, peptides were re-dissolved in 20 μl loading solvent A (0.1% TFA in water/ACN (98:2, v/v)) of which 5 μl was injected for LC-MS/MS analysis on an Ultimate 3000 RSLC nano LC (Thermo Fisher Scientific, Bremen, Germany) in-line connected to a Q Exactive mass spectrometer (Thermo Fisher Scientific). The peptides were first loaded on a trapping column that was made in-house, 100 μm internal diameter (I.D.) × 20 mm, 5 μm beads C18 Reprosil-HD (Dr. Maisch, Ammerbuch-Entringen, Germany). After flushing from the trapping column, the sample was loaded on an analytical column (made in-house, 75 μm I.D. × 150 mm, 5 μm beads C18 Reprosil-HD, Dr. Maisch) packed in the needle (PicoFrit SELF/P PicoTip emitter, PF360-75-15-N-5, New Objective, Woburn, MA, USA). Peptides were separated with a non-linear gradient from 98% solvent A’ (0.1% formic acid in water) to 30% solvent B′ (0.1% formic acid in water/acetonitrile, 20/80 (v/v)) in 21 min, reaching 56% solvent B′ at 29 min, at a flow rate of 250 nL/min. This was followed by a 10 min wash with 99% solvent B’ and re-equilibration with 98% solvent A’. The mass spectrometer was operated in data-dependent, automatically switching between MS and MS/MS acquisition for the 5 most abundant peaks in each MS spectrum. Full-scan MS spectra (400–2,000 m/z) were acquired at a resolution of 70,000 in the Orbitrap analyzer after accumulation to a target value of 3,000,000. The 5 most intense ions above a threshold value of 13,000 were isolated with a width of 2 m/z for fragmentation at a normalized collision energy of 25% after filling the trap at a target value of 50,000 for maximum 80 ms. MS/MS spectra (200–2000 m/z) were acquired at a resolution of 17,500 in the Orbitrap analyzer. The polydimethylcyclosiloxane background ion at 445.12002 Da was used for internal calibration (lock mass).

For PL, peptides were re-dissolved in 20 μl loading solvent A (0.1% TFA in water/ACN (98:2, v/v)) of which 1 μl was injected for LC-MS/MS analysis on an Ultimate 3000 RSLC nano LC (Thermo Fisher Scientific, Bremen, Germany) in-line connected to a Q Exactive mass spectrometer (Thermo Fisher Scientific). The peptides were first loaded on a PepMap^TM^ Neo Trap column (Thermo Fisher Scientific). After flushing from the trapping column, peptides were separated on a 50 cm μPAC™ column with C18-endcapped functionality (Pharmafluidics, Belgium) kept at a constant temperature of 50°C. Peptides were eluted by a non-linear gradient starting at 1% MS solvent B (0.1% FA in water/ACN, 20/80 (v/v)) and 99% MS solvent A (0.1% FA in water). For the gradient, MS solvent B reached 10% after 9 min, 33% at 32 min, 55% at 43 min, 70% at 48 min, followed by a 2 min wash with 70% MS solvent B and re-equilibration with 1% MS solvent B. Flow rate was set at 0.75 µL/min for 9 min and switched to 0.3 µL/min for the rest of the run. The mass spectrometer was operated in data-dependent mode, automatically switching between MS and MS/MS acquisition for the 5 most abundant peaks in each MS spectrum. Full-scan MS spectra (400–2,000 m/z) were acquired at a resolution of 70,000 in the Orbitrap analyzer after accumulation to a target value of 3,000,000 for maximum 80 ms. The 5 most intense ions above a threshold value of 13,000 were isolated with a width of 2 m/z for fragmentation at a normalized collision energy of 25% after filling the trap at a target value of 50,000 for maximum 80 ms. MS/MS spectra (200–2000 m/z) were acquired at a resolution of 17,500 in the Orbitrap analyzer. The polydimethylcyclosiloxane background ion at 445.12002 Da was used for internal calibration (lock mass).

### Proteomic data analysis and visualization

Protein identification was performed with the Mascot Distiller software (version 2.5.0.0 and 2.8.2.0, Matrix Science), in combination with the Mascot search engine (versions 2.5.1 and 2.8.2, Matrix Science), using the Mascot Daemon interface. Grouping of spectra was allowed with a max intermediate RT of 30 s and a max intermediate scan count of 5 was used where possible. Grouping was done with 0.005 Da precursor tolerance. A peak list was only generated when the MS/MS spectrum contained more than 10 peaks. There was no de-isotoping and the relative S/N limit was set to 2. Database was TAIR10plus^83^ for AP-MS, and Araport11plus^84^ for PL. Variable modifications were set to oxidation(M), acetyl(protein N-termini), and for PL experiments, also biotin(K). Mass tolerance on MS was set to 10 ppm (with Mascot’s C13 option set to 1) and the MS/MS tolerance at 20 mmu. The peptide charge was set to 2+, 3+ and 4+ and the instrument to ESI-QUAD. Trypsin was set as protease, allowing for 2 missed cleavages, and cleavage when R or K is followed by P. Only high-confident peptides, ranked 1 and with scores above the threshold score, set at 99% confidence, were withheld. Proteins with at least 1 unique peptide identified in at least 2 replicates were initially retained.

To determine bait-specific interactors in the AP-MS data, for all identifications, normalized spectral abundance factors (NSAF) in the 3 replicates were compared to the NSAFs in a large in-house dataset consisting of 809 AP-MS experiments performed with standard conditions. Thereto, the ratio of the average bait and control NSAF values were determined as a measure of fold enrichment, and a two-tailed Student’s t-test was performed comparing bait and control Ln-transformed NSAF values to determine the enrichment significance. t-test p-values were -Log10 transformed and infinite values were replaced by a constant -Log10 p-value of 325. Secondly, to correct for possible digitonin background, the identified proteins were compared to an in-house AP-MS dataset containing 131 experiments performed with 1% digitonin. Proteins were retained as specific if their Affinity Enrichment Score (AES), i.e. the product of their NSAF ratio and -Log10(p-value), in both comparisons was ≥ 20. Similarly, to determine bait-specific interactors in the PL data, a large dataset of 326 PL experiments were used for comparison. Proteins were retained as specific if their AES was ≥ 20.

Functional annotation of the interactomic data was performed using the ShinyGO web server (http://bioinformatics.sdstate.edu/go/)^85^ against the *Arabidopsis thaliana* reference genome database. Statistical significance for global interactome functional categories was evaluated using default parameters, with a False Discovery Rate (FDR) correction threshold set at P < 0.05. For the spatial proteomics sub-lists from the LOPIT datasets, functional enrichment profiling was conducted using the g:GOSt tool within the g:Profiler suite (https://biit.cs.ut.ee/gprofiler/)^86^. To maximize sensitivity in these smaller groups, a Benjamini-Hochberg FDR correction (*P* < 0.05) was applied in the analysis. Bubble plot representations were generated using GraphPad Prism 9 (GraphPad Software Inc.).

All protein-protein interaction (PPI) networks were constructed and visualized using Cytoscape software (v3.10.4)^87^. For the primary interactome network (**Fig. 1a**), node and label sizes were mapped proportionally to the log10 of a cumulative rank score, calculated by ranking proteins according to their enrichment values within each dataset and summing values for shared candidates to optimize visual scaling. Further refinements of the network were performed using Adobe Illustrator (Adobe Inc.).

### Protoplasts preparation and transfection

Protoplast isolation and transfection were performed using the tape-Arabidopsis sandwich method with some modifications^88^. In brief, abaxial epidermis of fully expanded 5-week-old *Arabidopsis* leaves grown under short-day-conditions was removed using tape, and submerged into enzymatic solution (20 mM MES (pH 5.7), 1.5% (w/v) cellulisine, 0.4% (w/v) macerozyme, 0.4 M mannitol, 20 mM KCl, 10 mM CaCl2, 1 mM beta-mercaptoethanol, and 0.1% (w/v) BSA) for protoplasts release. After 1.5 h room temperature incubation at 40 rpm, protoplasts were collected into round-bottom tubes and centrifuged at 100 g for 2 min. All centrifugations were performed with minimal acceleration and no brake to prevent cell lysis. The supernatant was discarded, and protoplasts were washed twice with W5 solution (2 mM MES pH 5.7, 154 mM NaCl, 125 mM CaCl2, and 5 mM KCl). After the last wash, protoplast suspension was incubated on ice for 30 min, pelleted and resuspended in MMG solution (4 mM MES pH 5.7, 0.4 M mannitol, and 15 mM MgCl2). Cell number was measured using a Neubauer chamber, and protoplast density was adjusted to approximately 4 x 10^5^ cells/mL with MMG solution.

Maxi-preps of plasmids were performed using the NucleoBond Xtra Maxi kit (Macherey-Nagel, Germany) and final plasmid concentration was adjusted to 1 μg/μL. For transfection, 100 μL of protoplasts suspension was transfected with 10 uL of plasmid and 100 μL of PEG solution (40% w/v PEG 4000, 0.2 M mannitol, and 100 mM CaCl_2_). After 10 minutes of incubation, transfection was stopped with 660 μL of W5 solution, followed by pelleting and resuspension in WI solution (4 mM MES pH 5.7, 0.5 M mannitol, 20 mM KCl, and 1 mM glucose). Transfected protoplasts were then transferred to 6-well plates, previously precoated with 5% BSA, and incubated a minimum of 18 hours at room temperature to allow protein expression. After incubation, protein expression and localization were checked on a confocal laser scanning microscope as described below (see Confocal microscopy section).

### Co-immunoprecipitation

For co-immunoprecipitation (CoIP), total proteins were extracted from approximately 10^5^ protoplasts per sample. Protein extraction was performed resuspending cell pellets in 500 μL of cold Extraction buffer (150 mM Tris-HCl (pH 7.5), 150 mM NaCl, 10% (v/v) glycerol, 10 mM EDTA, 1 mM NaF, 1 mM Na-molybdate, and 0.2% (v/v) Nonidet P-40, 10 mM DTT, 0.5 mM PMSF, 1% (v/v) protease inhibitor P9599 (Sigma, USA)).

Suspensions were then incubated for 10 min at 4°C on an end-over-end rotator and centrifuged at 15000g for 20 min at 4°C. Supernatants were transferred to low-binding tubes and 50 μL of sample was reserved as input. The remaining volume was incubated 2 h at 4°C with 25 uL of GFP-Trap Magnetic Agarose beads, (Chromotek, Germany), previously washed three times with 500 μL of Wash buffer (same composition as Extraction buffer with 0.05% NP-40). After incubation, beads were washed three times with 500 μL of Wash buffer, changing to a new tube during the last washing step to minimize carryover. After last wash, beads were suspended in 50 uL of 2X Laemmli buffer (125 mM Tris-HCl pH 6.8, 4% SDS, 20% glycerol, 0.01% bromophenol blue) supplemented with 2% (v/v) beta-mercaptoethanol. Input samples were also mixed with equal volume of 4X Laemmli buffer supplemented with 4% (v/v) beta-mercaptoethanol. Protein denaturation was performed by heating at 70 °C for 30 min with frequent vortexing, and samples were centrifuged 10 min at 20000g prior to Western blot analysis, as described below.

### Western Blot

For immunoblot analysis, input and CoIP samples were separated by SDS-PAGE using Mini-PROTEAN TGX Precast Gels (Bio-Rad, USA) and transferred onto Immobilon-P 0.45 μm PVDF membranes (Sigma) using the Trans-Blot Turbo Transfer System (Bio-Rad). Membranes were blocked for 2 h with 5% milk in TTBS and then incubated overnight at 4 °C in TTBS with mouse monoclonal anti-GFP 1:600 (sc-9996, Santa Cruz Biotechnology, USA) or rabbit polyclonal anti-SYT1 1:1000. Membranes were then washed three times with TTBS and incubated 1 h at room temperature with the HRP-conjugated antibodies anti-mouse 1:80000 (A9044, Sigma) or anti-rabbit 1:80000 (A0545, Sigma). After three more washes, protein detection was performed using either Clarity Western ECL (Bio-Rad) or SuperSignal West Atto (Thermo Fisher Scientific). Uniform protein loading was assessed staining with Coomasie Brilliant Blue R-250 (CBB).

### Nicotiana benthamiana transient expression

For transient expression in *N. benthamiana*, the respective constructs were transformed into *Agrobacterium tumefaciens* GV3101 (pMP90) and selected on LB supplemented with 50 µg/mL rifampicin, 25 µg/mL gentamicin and 100 µg/mL spectinomycin / 50 µg/mL kanamycin for plasmid selection. Bacterial cultures were harvested by centrifugation at 3000g for 15 min and resuspended in agroinfiltration solution (10 mM MES pH 5.6, 10 mM MgCl2, and 1 mM acetosyringone). Suspensions were then incubated for 2 h in the dark at room temperature. Prior to infiltration, cultures were mixed to achieve specific optical densities at 600 nm (OD600), always including the p19 silencing suppressor. For single expression, OD600 values were adjusted to 0.7 for the target construct and 0.25 for p19. For double expression assays (including co-expression, BiFC, and Sac experiments), OD600 values were set to 0.3 for each target construct and 0.15 for p19. For triple expression, target constructs were mixed at an OD600 of 0.2 each, with p19 at 0.15. Fully expanded leaves of 3-week-old *N. benthamiana* were syringe-infiltrated on the abaxial side, and confocal imaging was performed 2 days after infiltration as described below.

### Isoxaben treatment and lignin staining

For cell wall stress assays, 5-day-old vertically grown *Arabidopsis* seedlings (Col-0, *syt1*, and *pSYT1:SYT1-GFP*) were treated with the cellulose biosynthesis inhibitor isoxaben (ranging from 6 to 600 µM) or mock-treated for 18 h in half-strength liquid MS medium. For ectopic lignin visualization, seedlings were fixed in an ethanol to acetic acid solution (3:1, v/v) for 2 h and rehydrated through a graded ethanol series (75%, 50%, and 25% ethanol) for 30 min per step. After an overnight incubation in distilled water, fixed seedlings were histochemically stained using 2:1 concentrated hydrochloric acid to 1% phloroglucinol. Stained samples were immediately mounted in water and imaged. Images were captured using a Zeiss SteREO Discovery.V12 stereomicroscope coupled to an AxioCam 503 digital camera.

### Confocal imaging

Microscopy images were obtained using a Zeiss LSM 880 confocal laser scanning microscope (Carl Zeiss, Germany). Sample images were taken using a water-immersion C-Apochromat 40x/1.2 W Korr objective or Plan-Apochromat 63x/1.40 Oil DIC immersion objective, with up to 3x zoom. For imaging in *N. benthamiana*, leaf-discs were mounted in water and transient expression was observed on abaxial epidermis cells. For imaging of transfected protoplasts, 10 µL of resuspended protoplast suspension was mounted directly onto a Neubauer chamber for visualization. For *Arabidopsis* root imaging, seedlings were mounted in half-strength liquid MS medium, imaging cells from the transition zone at approximately the same distance from the quiescent center. When required, plasma membrane staining was done by incubating seedlings 5 min with 2 µM FM4-64 (Thermo Fisher Scientific). Detection was performed using PMT or GaAsP detectors, and an additional PMT for transmitted light. Excitation wavelengths and emission detection windows were optimized for each target as follows: a 405 nm diode laser was used to excite EBFP (Em. 410-485 nm) and ectopic lignin autofluorescence (Em. 510-580 nm); a 488 nm Argon laser was used for EGFP (Em. 499-590 nm), reconstituted EYFP (Em. 510-620 nm), and FM4-64 (Em. 620-750 nm); and a 561 nm DPSS laser was employed for mCherry (Em 568-658 nm). Images were obtained as single planes or Z-stacks from the equatorial plane to the cell cortex. Microscopy images were processed using Fiji^89^. When required, image drift was corrected using the StackReg plugin^90^.

### Image segmentation and quantification

Semiautomatic image segmentation was performed using the machine learning-based software Ilastik^62^. For membrane contact site (MCS) detection of transiently expressed proteins in *N. benthamiana*, the pixel classifier was trained using cortical images to distinguish discrete puncta, reticular ER strands, background, and overexposed regions. Quantification was performed by calculating the surface area ratio between punctate and reticulated signals from the generated binary masks. For lignin quantification in *Arabidopsis* roots, the pixel classifier was trained using stereomicroscope images of mock- and isoxaben-treated seedling roots to separate phloroglucinol-stained areas from non-lignified root tissues and background signals. Quantification was performed by extracting the absolute values (in pixels) of the lignified surface area from the segmented images.

### Statistical analysis and illustrations

All graphs and statistical analyses were performed using GraphPad Prism 9. Quantitative data are presented as violin plots displaying individual data points, with horizontal lines indicating the mean and quartiles. For multi-group comparisons, statistical significance among genotypes within each individual group was determined using a Brown-Forsythe ANOVA followed by a Games-Howell *post-hoc* test to account for unequal variances. Statistical significance was defined as *P* < 0.001, with significant differences indicated by letters or asterisks in the figures. Schematic models and diagrams were created using BioRender.com and Adobe Illustrator.

## ACKNOWLEDGMENTS

We would like to thank Dominique Eeckhout and the ViB Interactomics Facility for the LC-MS/MS analyses; Wout Boerjan, Raquel Pagano and Ángel del Espino for their insightful discussion and feedback of the project; and Alicia Esteban del Valle for technical assistance with confocal imaging.

This work was supported by the Ministry of Science and Innovation, grant no. 775 (PID2023-147983OB-I00), awarded to M.A.B and L.R.; PPRO-AGR168-G-2023PPRO-AGR168-G-2023 project by Junta de Andalucía, awarded to M.A.B.; the European Research Council (ERC) grant agreement no. 815 (101001097-LIPIDEV), awarded to Y.J.; Programa Emergia 2023 grant (DGP_EMEC_2023_00375) from Junta de Andalucía, awarded to V.A-S; and Ramón y Cajal contract RYC2024-049082-I from the Spanish Ministerio de Ciencia, Innovación y Universidades, awarded to J.P-S. J.M-L. was funded by Researcher Training Fellowship PRE2021-097655. U.G was funded by Researcher Training Fellowship PRE2023-001276. V.M. was funded by European Molecular Biology Organization (EMBO) fellowship ATLF 466-2022.

## AUTHOR CONTRIBUTIONS

J.M-L. and M.A.B. wrote the manuscript. J.M-L., M.A.B. and Y.J. conceived the research. J.M-L., U.G., F.B-F., V.M., J.P-S. and V.A-S. performed experiments. J.V.L. and G.D.J. generated and analyzed the mass spectrometry data. H.P. and T.J.S. performed the LOPIT dual-localization analyses. J.M-L. and U.G. generated data visualizations and figures. M.A.B., Y.J. and L.R. supervised the project. M.A.B., L.R., Y.J., V.A-S. and J.P-S. secured funding. All authors contributed to manuscript revision and approved the final version of the text.

## COMPETING INTERESTS

The authors declare no competing interests.

## Notes

### Competing Interest Statement

The authors have declared no competing interest.

