## Supplementary Figures and Tables for "Integration of SYT1 Interactomics and Dual-Localization Proteomics Links ER-PM Contacts to Lignin Deposition"

**Supplementary Figure 1.** Expanded SYT1 protein-protein interaction network.

**Supplementary Figure 2.** Gene Ontology Molecular Function enrichment analysis of the global SYT1 interactome.

**Supplementary Figure 3.** LOPIT ER-PM dual-localization thresholds optimization.

**Supplementary Figure 4.** Subcellular localization of SEPC1 and SEPC2 fusions in *Arabidopsis* protoplasts.

**Supplementary Figure 5.** Comprehensive metabolic map of the monolignol biosynthetic pathway leading to lignin precursors.

**Supplementary Figure 6.** Subcellular localization of transiently expressed MSBP1 and MSBP2 fusions.

**Supplementary Figure 7.** Machine-learning based quantification of root ectopic lignification.

**Supplementary Figure 8.** Dose-dependent assay of isoxaben-induced ectopic lignification in Col0, *syt1* and SYT1-GFP complemented lines.

**Supplementary Figure 9.** Proposed models for monolignols export.

**Supplementary Video 1.** Co-accumulation of MSBP2 at immobile SYT1-positive cortical foci.

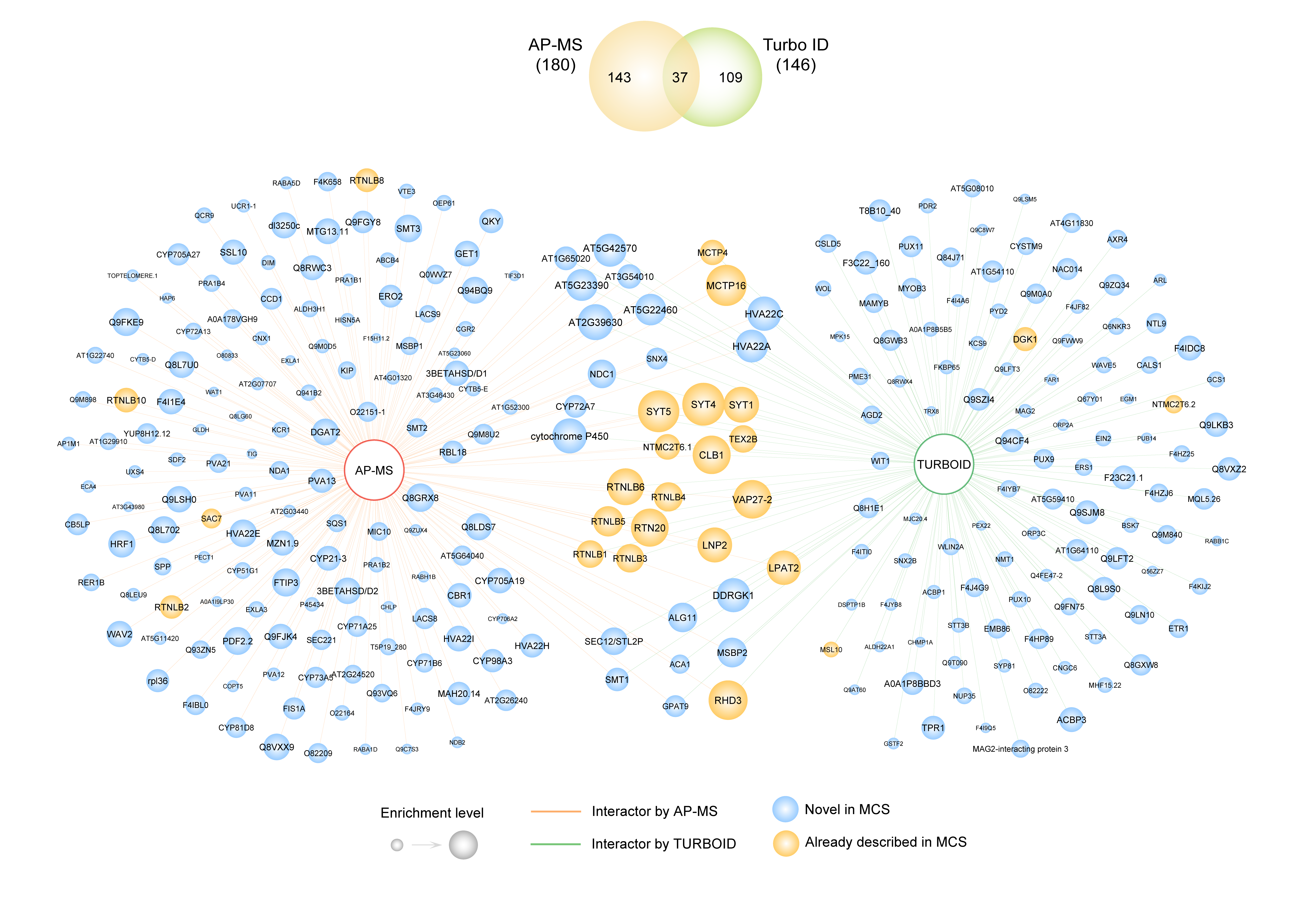

#### **Supplementary Figure 1. Expanded SYT1 protein-protein interaction network.**

Node and edge attributes are identical to those described in Figure 1a, but with all nodes labelled. The Venn diagram (top) illustrates the overlap of proteins across the AP-MS and PL experiments.

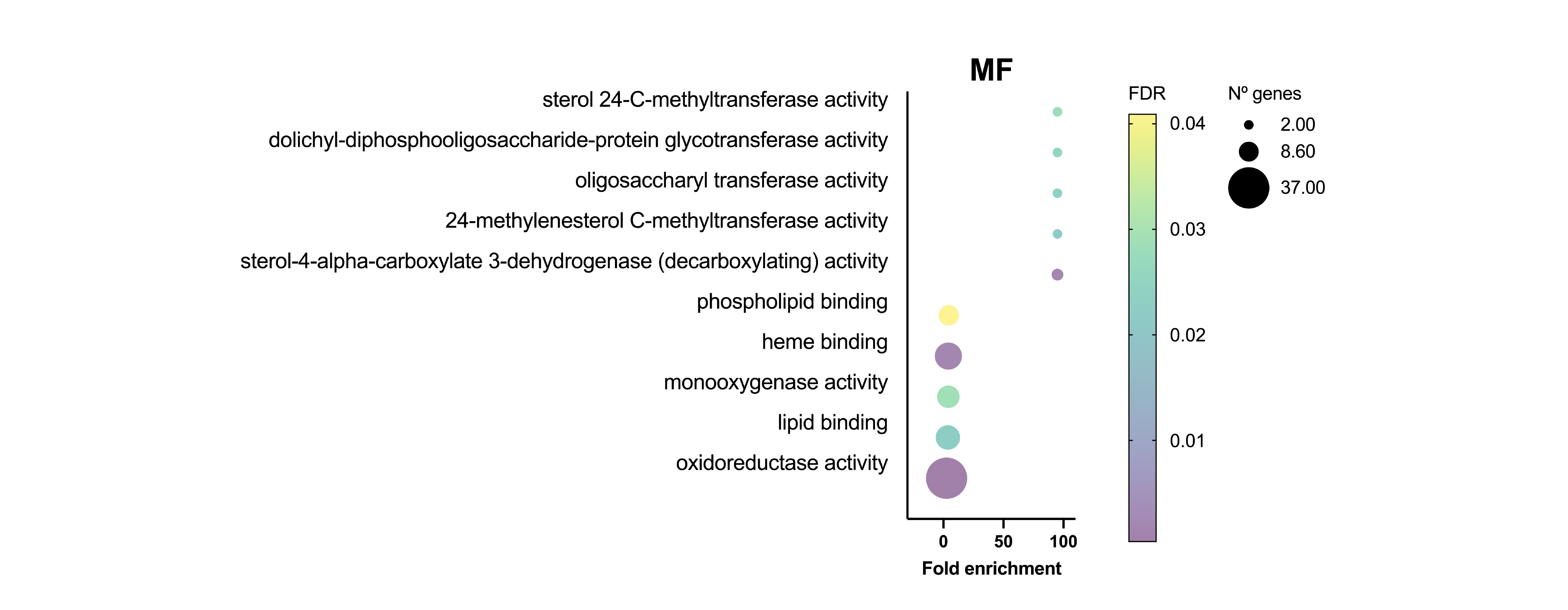

#### Supplementary Figure 2. Gene Ontology Molecular Function enrichment analysis of the global SYT1 interactome.

Dot plot displaying significantly enriched Molecular Function (MF) terms for the combined AP-MS and PL datasets. The x-axis indicates fold enrichment, the bubble size corresponds to the number of genes assigned to each term, and the color gradient represents the false discovery rate (FDR) significance level. Annotated terms highlight the expected prominence of lipid-binding and steroid-related pathways alongside less anticipated activities.

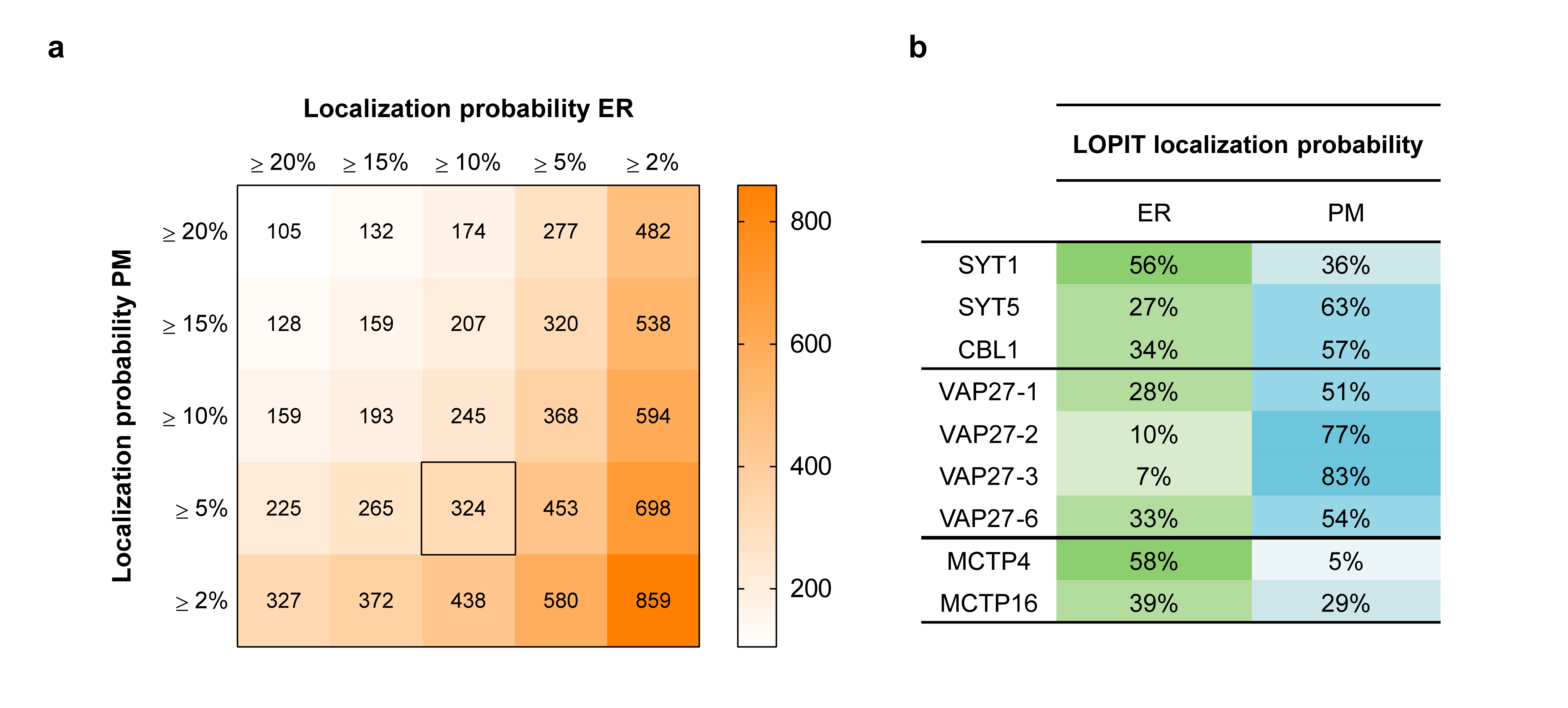

#### Supplementary Figure 3. LOPIT ER-PM dual-localization thresholds optimization.

**a, Heatmap displaying the total number of candidate proteins recovered across varying LOPIT assignment probability thresholds.** Endoplasmic Reticulum (ER) probabilities are plotted on the x-axis, while Plasma Membrane (PM) probabilities lie on the y-axis. The black square outlines the selected thresholds for ER-PM dual localization (≥ 10% ER, ≥ 5% PM), yielding 324 ER-PM dual-localized proteins. **b, LOPIT localization probabilities (%) for canonical ER-PM contact sites components.** Representative members of the Synaptotagmin, VAP27, and MCTP families are shown.

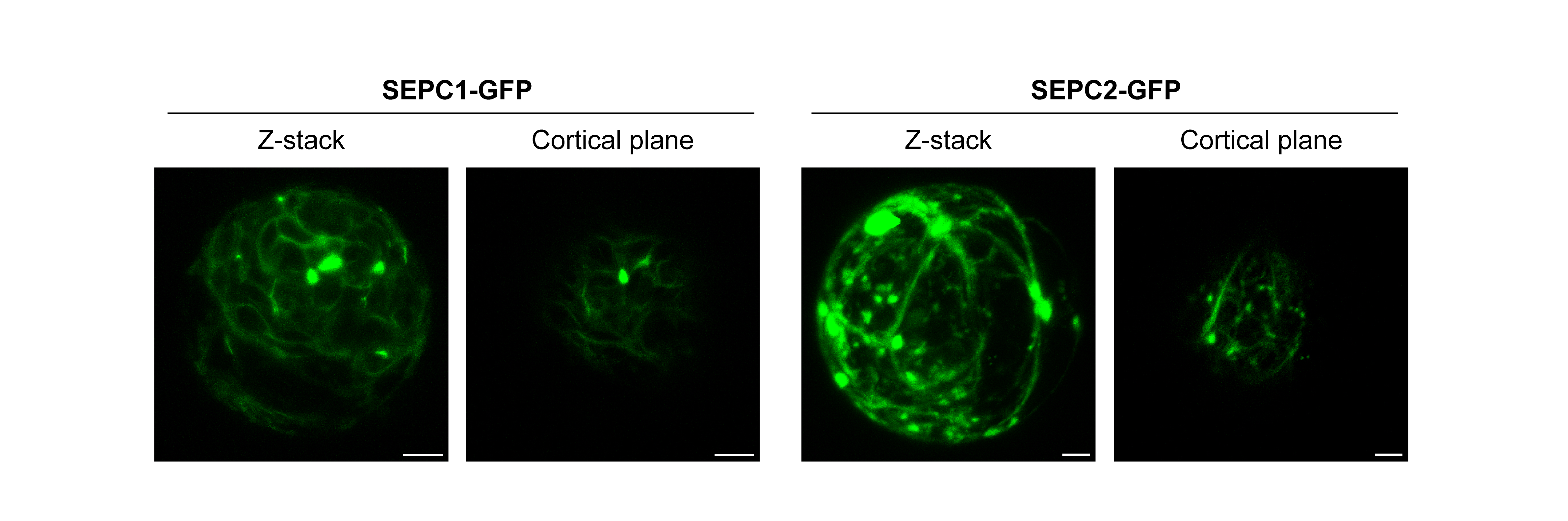

#### Supplementary Figure 4. Subcellular localization of SEPC1 and SEPC2 fusions in *Arabidopsis* protoplasts.

Confocal images displaying the subcellular distribution of transiently expressed SEPC1-GFP and SEPC2-GFP in *Arabidopsis* protoplasts prior to co-immunoprecipitation assays. Maximum intensity projections of full Z-stacks (left) and more detailed cortical sections (right) are shown for each protein. Scale bars, 5 µm.

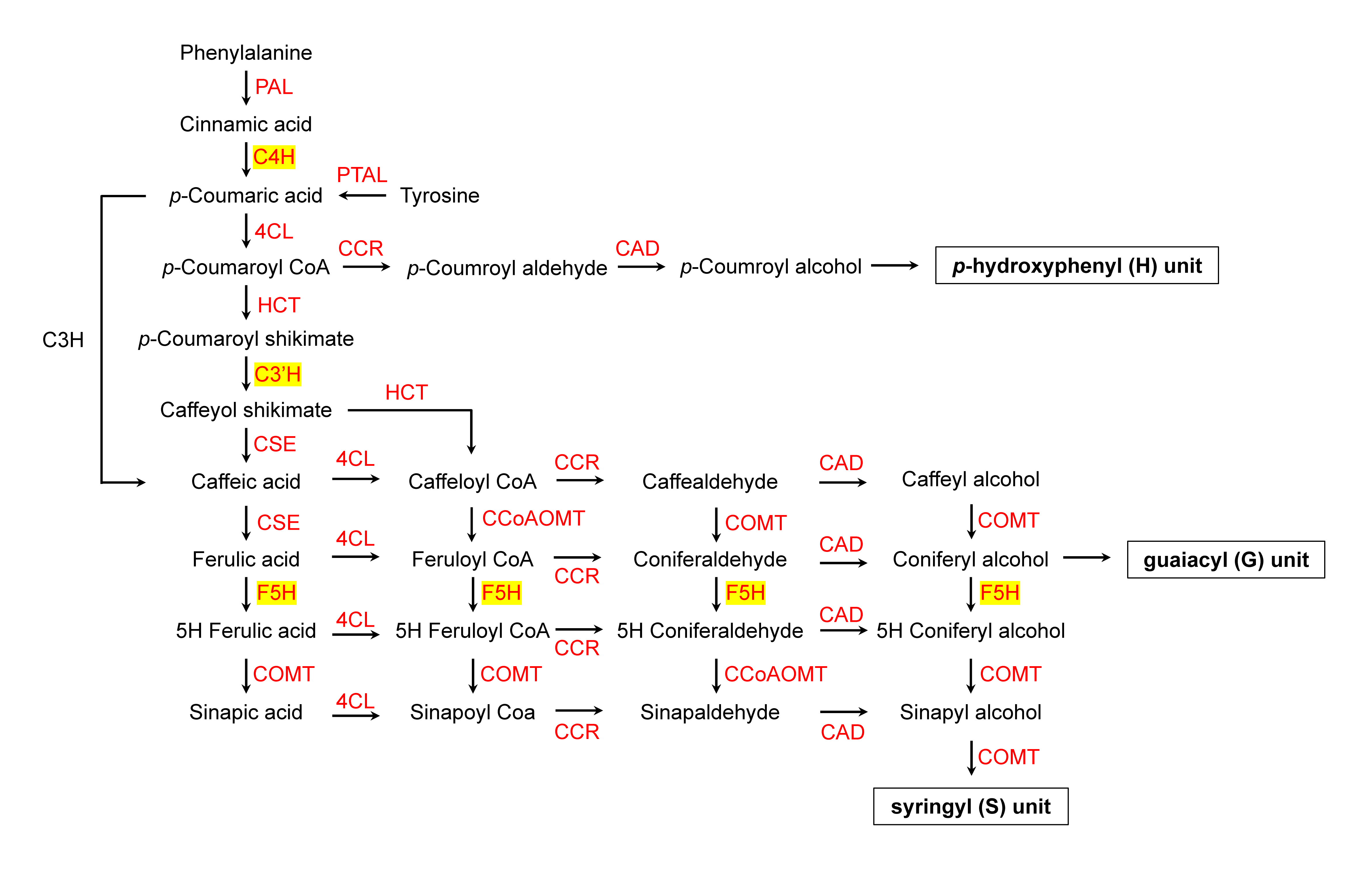

#### Supplementary Figure 5. Comprehensive metabolic map of the monolignol biosynthetic pathway leading to lignin precursors.

Diagram illustrating the sequential enzymatic steps, chemical intermediates, and metabolic branches of the phenylpropanoid pathway leading to the synthesis of p-hydroxyphenyl (H), guaiacyl (G), and syringyl (S) lignin subunits in plants. Core cytochrome P450 monooxygenases investigated in this study (C4H, C3'H, and F5H) are highlighted in yellow boxes. Associated catalytic enzymes responsible for each reaction step are indicated in red. Abbreviations: PAL, phenylalanine ammonia-lyase; PTAL, phenylalanine/tyrosine ammonia-lyase; C4H, cinnamate 4-hydroxylase; 4CL, 4-coumarate:CoA ligase; HCT, hydroxycinnamoyl-CoA:shikimate hydroxycinnamoyltransferase; C3'H, p-coumarate 3'-hydroxylase; CSE, caffeoyl shikimate esterase; CCoAOMT, caffeoyl-CoA O-methyltransferase; F5H, ferulate 5-hydroxylase; COMT, caffeic acid O-methyltransferase; CCR, cinnamoyl-CoA reductase; CAD, cinnamyl alcohol dehydrogenase; H, p-hydroxyphenyl unit; G, guaiacyl unit; S, syringyl unit.

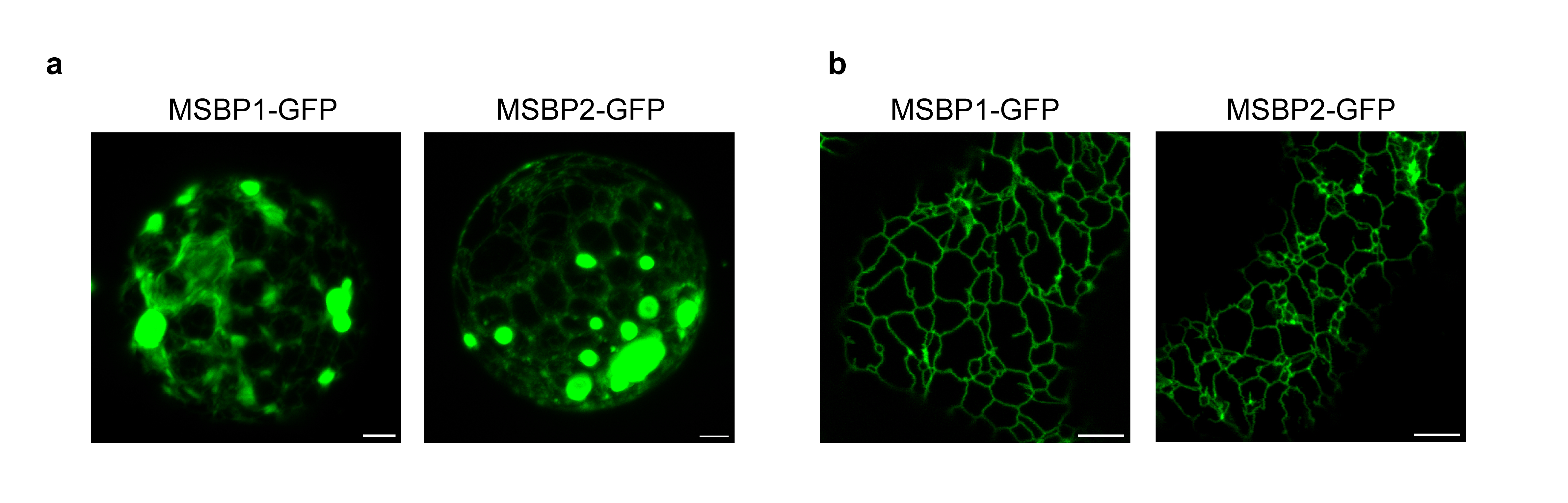

#### Supplementary Figure 6. Subcellular localization of transiently expressed MSBP1 and MSBP2 fusions.

**a, Subcellular localization of MSBP1 and MSBP2 in *Arabidopsis* protoplasts.** Confocal micrographs displaying the distribution of transiently expressed MSBP1-GFP and MSBP2-GFP fusions in *Arabidopsis thaliana* mesophyll protoplasts prior to co-immunoprecipitation assays. **b, Cortical ER distribution of MSBP1 and MSBP2 in *Nicotiana benthamiana*.** Confocal images showing a cortical view of epidermal cells transiently expressing MSBP1-GFP or MSBP2-GFP. Scale bars, 5 µm.

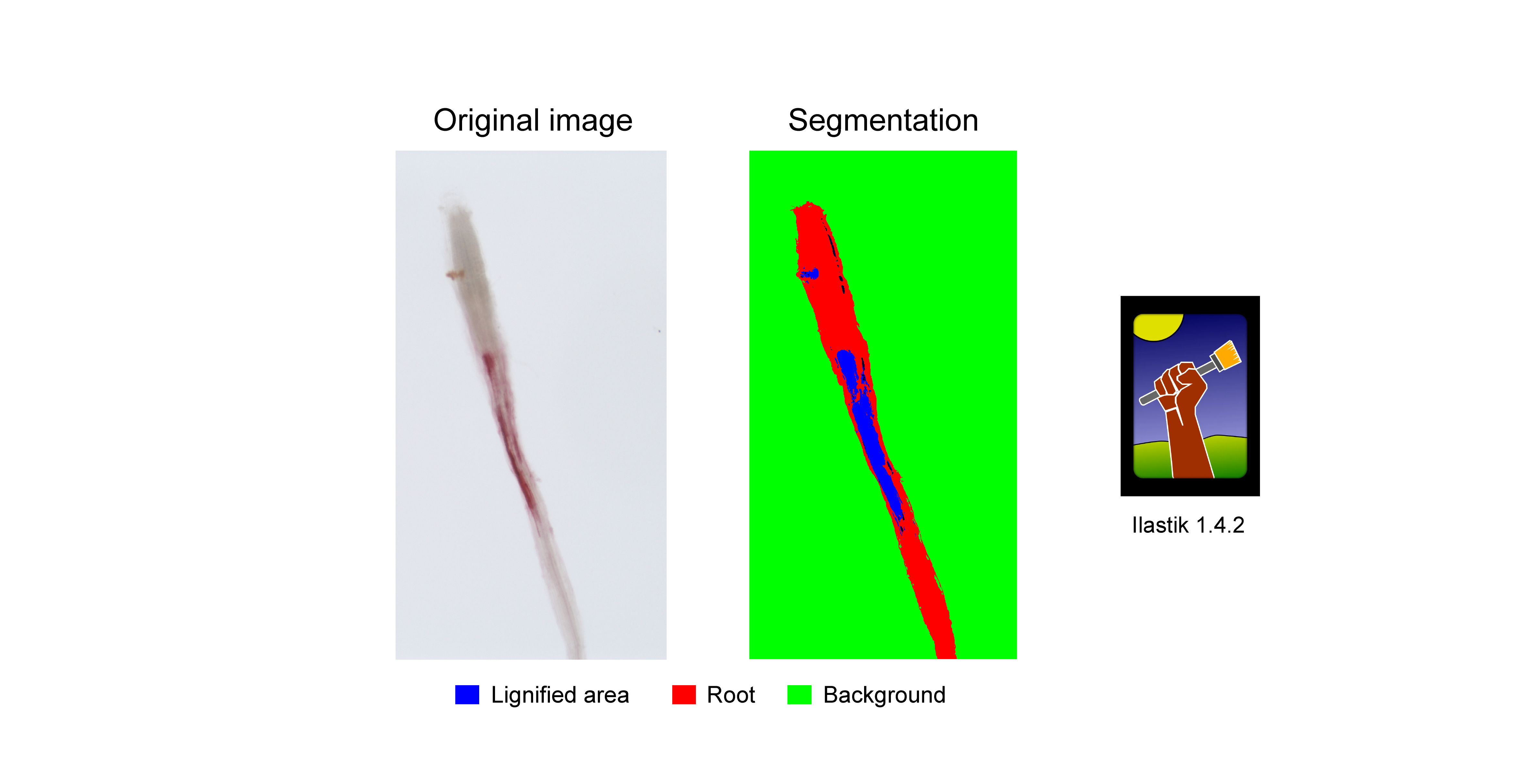

#### Supplementary Figure 7. Machine-learning based quantification of root ectopic lignification.

Automated pixel classification strategy used for the unbiased quantification of phloroglucinol-HCl-stained roots. Unbiased segmentation was performed using the machine-learning and segmentation toolkit Ilastik 1.4.2.

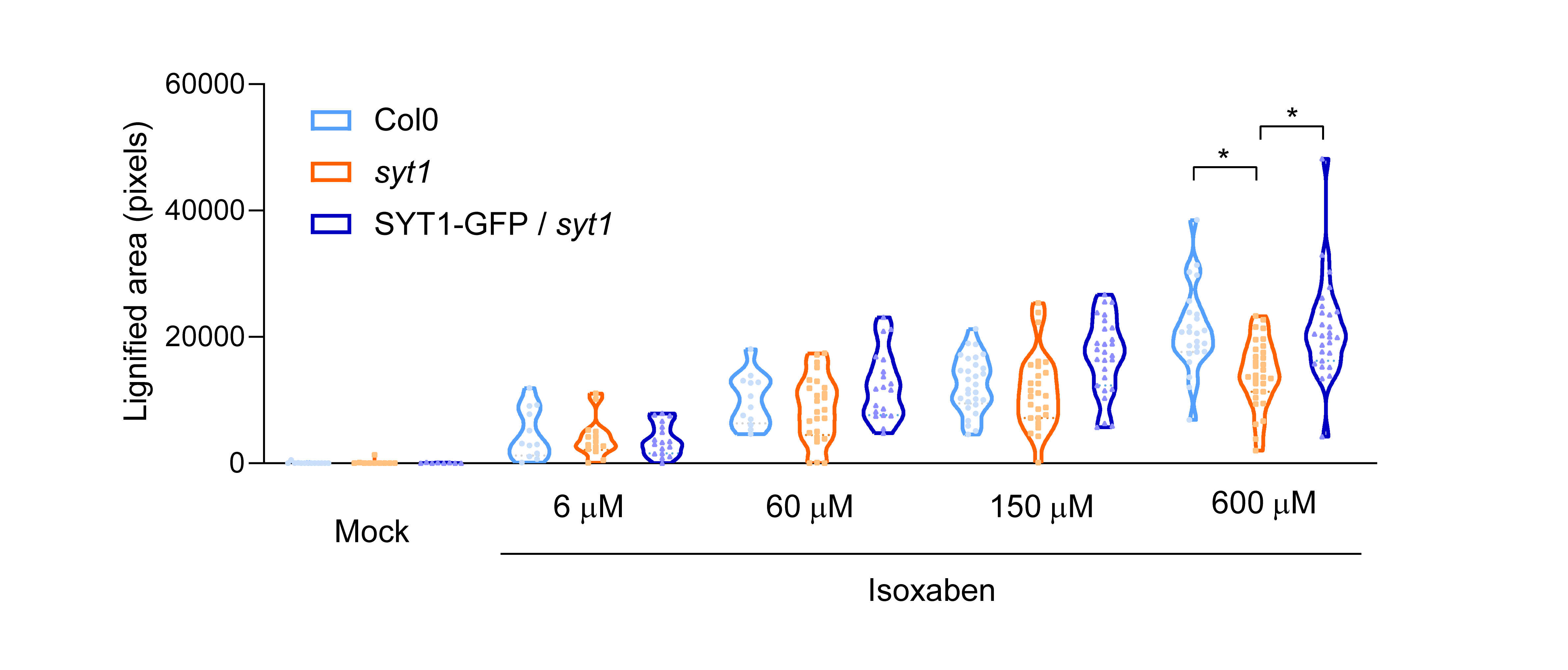

#### Supplementary Figure 8. Dose-dependent assay of isoxaben-induced ectopic lignification in Col0, *syt1* and SYT1-GFP complemented lines.

Violin plots displaying the quantification of the ectopic lignified root area across an 18 h treatment with different concentrations (0, 6, 60, 180 and 600 µM) of the cellulose biosynthesis inhibitor isoxaben. Data are presented as violin plots with individual data points; horizontal lines indicate the mean and quartiles (n ≥ 10 images). Statistical analysis was performed among genotypes within each individual group. Statistical significance was determined by a Brown-Forsythe ANOVA followed by a Games-Howell post-hoc test for multiple comparisons. Single asterisk (*) denote pairs with significant differences (*P* < 0.001).

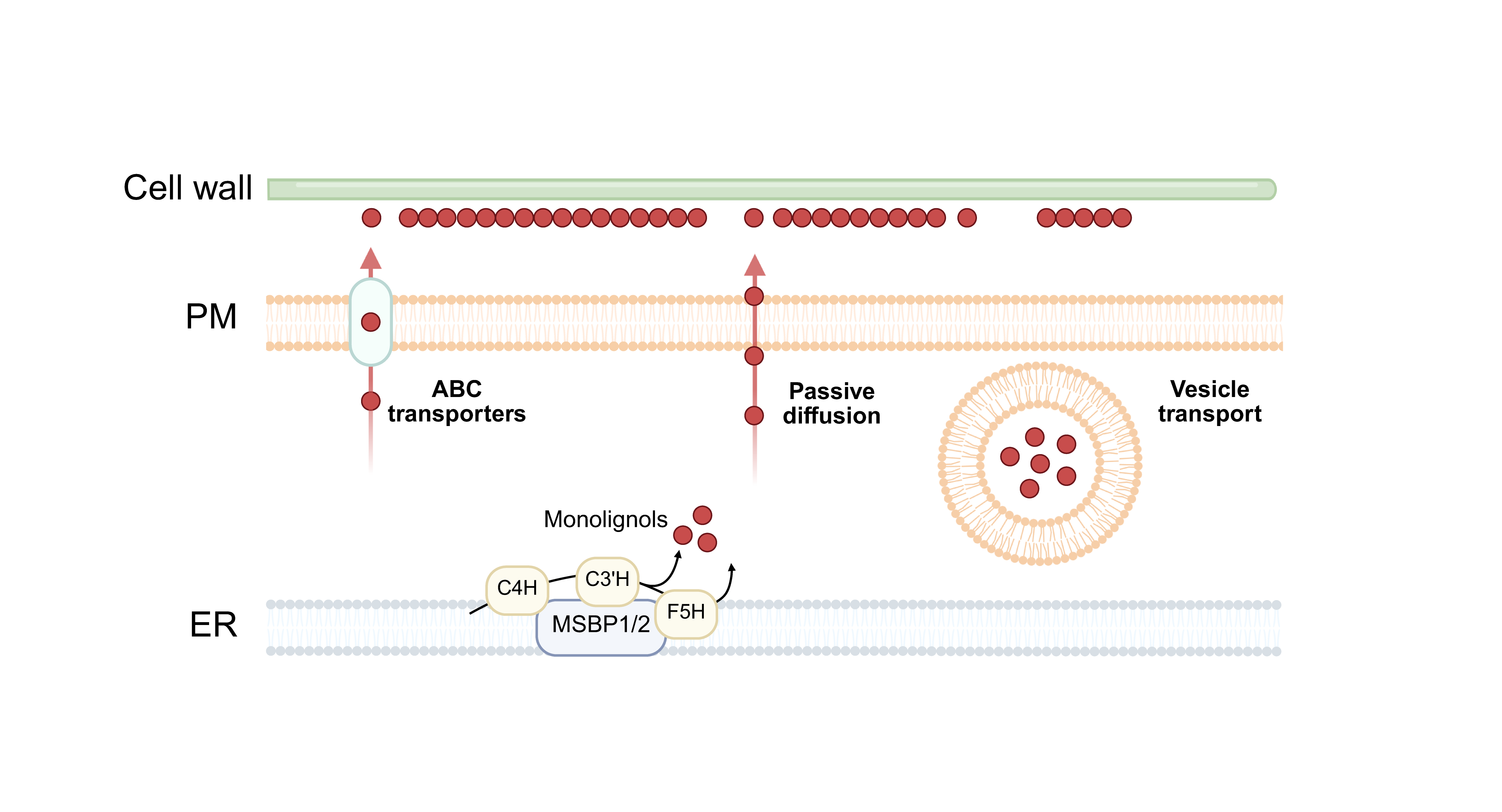

#### Supplementary Figure 9. Proposed models for monolignols export.

Schematic overview depicting the three primary proposed mechanisms for monolignol transport from the endoplasmic reticulum (ER) to the cell wall: ABC-type transporters, passive bilayer diffusion, and vesicle-mediated trafficking.

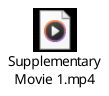

#### Supplementary Video 1. Co-accumulation of MSBP2 at immobile SYT1-positive cortical foci.

Time-lapse confocal imaging of *Nicotiana benthamiana* epidermal cells transiently co-expressing MSBP2-GFP (left, green) and SYT1-mCherry (center, magenta). The merged channels (right) demonstrate the accumulation of MSBP2 into discrete, SYT1-positive cortical microdomains. White arrowheads highlight immobile puncta. Total acquisition time course, 54.8 s. Scale bars, 5 µm.

### SUPPLEMENTARY TABLES

**Supplementary Table 1.** Identified SYT1 interactors related to membrane contact site biology and endoplasmic reticulum architecture.

**Supplementary Table 2.** Cross-reference of SYT1 interactors against already published CYP73A5/CYP98A3 and RTNLB3/RTNLB6 proteomes.

#### Supplementary Table 1. Identified SYT1 interactors related to membrane contact site biology and ER architecture.

| **Protein name** | **AGI Identifier** | **Relation with MCS** | **Reference** |
| --- | --- | --- | --- |
| CLB1 | AT3G61050 | SMP-domain family SYT1 interactor (targeted assay) | ^33^ |
| DGK1 | AT5G07920 | SYT1 interactor (targeted assay) | ^18^ |
| LNP2 | AT4G31080 | ER-shaping protein | ^36^ |
| MCTP15 | AT1G74720 | MCTPs family | ^91^ |
| MCTP16 | AT5G17980 | MCTPs family | ^91^ |
| MCTP3 | AT3G57880 | MCTPs family | ^91^ |
| MCTP4 | AT1G51570 | MCTPs family | ^91^ |
| MSL10 | AT5G12080 | VAP27 interactor (targeted assay) | ^92^ |
| NTMC2T6.1 | AT1G53590 | SMP-domain family | ^93^ |
| NTMC2T6.2 | AT3G14590 | SMP-domain family | ^93^ |
| ORP2A | AT4G22540 | VAP27 interactor (targeted assay) | ^49^ |
| RHD3 | AT3G13870 | ER-shaping protein | ^68^ |
| RTN20 | AT2G43420 | Reticulon family | ^91^ |
| RTNLB1 | AT4G23630 | Reticulon family | ^91^ |
| RTNLB10 | AT2G15280 | Reticulon family | ^91^ |
| RTNLB11 | AT3G19460 | Reticulon family | ^91^ |
| RTNLB2 | AT4G11220 | Reticulon family | ^91^ |
| RTNLB3 | AT1G64090 | Reticulon family | ^91^ |
| RTNLB4 | AT5G41600 | Reticulon family | ^91^ |
| RTNLB5 | AT2G46170 | Reticulon family | ^91^ |
| RTNLB6 | AT3G61560 | Reticulon family | ^91^ |
| RTNLB8 | AT3G10260 | Reticulon family | ^91^ |
| SAC7 | AT3G51460 | SYT1 interactor (targeted assay) | See companion manuscript – Marković et al., 2026 |
| SYT1 | AT2G20990 | Bait | - |
| SYT4 | AT5G11100 | SMP-domain family | ^94^ |
| SYT5 | AT1G05500 | SMP-domain family SYT1 interactor (targeted assay) | ^95^ |
| TEX2B | AT1G73200 | SMP-domain family | ^93^ |
| VAP27-1 | AT3G60600 | VAP27 family | ^27^ |
| VAP27-2 | AT1G08820 | VAP27 family | ^27^ |
| VAP27-3 | AT2G45140 | VAP27 family | ^27^ |
| VAP27-4 | AT5G47180 | VAP27 family | ^27^ |
| VAP27-6 | AT4G00170 | VAP27 family | ^27^ |

#### Supplementary Table 2. Cross-reference of SYT1 interactors against already published CYP73A5/CYP98A3 proteome^76^.

| **AGI Identifier** | **AP-MS interactors of CYP73A5 (C4H) and CYP98A3 (C3’H)^90^** | **Presence in the SYT1 dataset** | **Identification method** |
| --- | --- | --- | --- |
| AT2G20990 | SYNAPTOTAGMIN 1 (SYT1) | YES | Bait |
| AT2G30490 | CYP73A5 (C4H) | YES | AP-MS |
| AT3G26830 | PAD3 | NO | - |
| AT5G22460 | SEPC1 | YES | AP-MS / TurboID / ER-PM LOPIT |
| AT4G22710 | CYP706A2 | YES | AP-MS |
| AT5G53560 | Cytochrome b5 isoform 1 | YES | AP-MS |
| AT4G23630 | Reticulon family protein (RTNLB1) | YES | AP-MS / TurboID / ER-PM LOPIT |
| AT4G11220 | Reticulon family protein (RTNLB2) | YES | AP-MS / ER-PM LOPIT |
| AT1G64090 | Reticulon family protein (RTNLB3) | YES | AP-MS / TurboID / ER-PM LOPIT |
| AT5G41600 | Reticulon family protein (RTNLB4) | YES | AP-MS / TurboID / ER-PM LOPIT |
| AT2G46170 | Reticulon family protein (RTNLB5) | YES | AP-MS / TurboID / ER-PM LOPIT |
| AT2G03510 | Band 7 family protein | NO | - |
| AT1G51570 | C2 domain-containing protein (MCTP4) | YES | AP-MS / TurboID |
| AT1G20330 | SMT2-sterol methyltransferase 2 | YES | AP-MS |
| AT5G52240 | MSBP1 | YES | AP-MS / ER-PM LOPIT |
| AT3G48890 | MSBP2 | YES | AP-MS / Turbo ID / ER-PM LOPIT |
| AT2G45140 | VAP27-3 | YES | AP-MS |
